# PKM2 diverts glycolytic flux in dependence on mitochondrial one-carbon cycle

**DOI:** 10.1101/2023.01.23.525168

**Authors:** Mohaned Benzarti, Anais Oudin, Elodie Viry, Ernesto Gargiulo, Maryse Schmoetten, Laura Neises, Coralie Pulido, Nadia I. Lorenz, Michael W. Ronellenfitsch, David Sumpton, Marc Warmoes, Christian Jaeger, Antoine Lesur, Etienne Moussay, Jerome Paggetti, Simone P. Niclou, Elisabeth Letellier, Johannes Meiser

## Abstract

Throughout the metastatic cascade, cancer cells are faced with harsh metabolic environments and nutritional stresses which apply selection pressure leaving only the most metabolically resilient cells to survive and form metastases. Metabolic characterisation of such cell populations *in vitro* is currently challenging. Using galactose as a tool compound to mimic glycolytic limitation within the tumour microenvironment of primary and secondary neoplastic sites, we were able to uncover metabolic flexibility and plasticity of cancer cells *in vitro*. In contrast to the established idea that high glycolytic flux and expression of dimeric PKM2 redirects carbons towards anabolic routes such as the pentose phosphate pathway and serine synthesis pathway (SSP), we have discovered by using stable-isotope tracing that also glycolytic limitation results in metabolic rewiring. Surprisingly, despite limited carbon availability and energetic stress, cells induce a near complete block of pyruvate kinase isozyme M2 (PKM2) to divert carbons towards SSP. Simultaneously, TCA cycle flux is sustained and oxygen consumption is increased, both supported by glutamine. Glutamine not only supports TCA cycle flux but also SSP via distinct mechanisms. Due to PKM2 block, malic enzyme exclusively supports TCA cycle flux while mitochondrial phosphoenolpyruvate carboxykinase supports SSP. Moreover, by using genetic modifications of different one-carbon (1C) cycle enzymes, we are able to reverse the PKM2 block suggesting a link between mitochondrial 1C cycle and pyruvate kinase. Thus we show that PKM2 inhibition acts as a branching point to direct glycolytic and glutamine carbons into distinct routes, overall supporting the metabolic plasticity and flexibility of cancer cells.

## Introduction

Increased aerobic glycolysis has been identified as a characteristic of cancer cells almost 100 years ago and subsequently highlighted as a hallmark of cancer [1-3]. This is the phenomenon whereby cancer cells increase the fermentation of glucose to lactate even in the presence of oxygen, at the cost of releasing and thereby losing carbons as lactate [4]. Many theories have been suggested as to why cancer cells adopt such a biomass-wasteful and energy-inefficient phenotype with the majority of these theories circulating around the idea that mitochondria are unable to cope with very high rates of early glycolytic reactions that are driven by oncogenic signalling [5-10]. In these early steps of glycolysis, NAD^+^ and ATP are consumed and need to be regenerated for glycolysis to continue. This regeneration of NAD^+^ and ATP seems to be the decisive bottleneck where the cancer cell has to decide whether to ferment glucose or oxidise it fully in the TCA cycle [9, 11-13]. Indeed, two recent publications highlighted how aerobic glycolysis rates are decided based on NADH shuttle capacity and the regeneration of NAD^+^ [12, 13]. Moreover, it was proposed that cells downregulate mitochondrial respiration to avoid ATP depletion when the mitochondria are unable to replenish ATP used in the early steps of glycolysis fast enough. Therefore, cells turn to ATP generation via lactate overflow to avoid an ATP crisis [4, 9, 11].

While it has been suggested that aerobic glycolysis is not the main driver of high proliferation rates [13], it remains undeniable that cancer cells engage high proliferative rates in favourable metabolic environments with ampleness of glucose being an example of such environments. Therefore, *in vitro* it might seem that cancer cells are fixed on aerobic glycolysis, while *in vivo* they actually display metabolic flexibility and plasticity [14]. This is to survive nutrient depleted and hostile metabolic environments within the local tumour microenvironment (TME) and along the metastatic cascade [15]. However, such metabolic flexibility is usually masked within *in vitro* studies due to the usage of synthetic media containing supraphysiological concentrations of various metabolites [16]. Aerobic glycolysis can be caused *in vitro* as cancer cells become reversibly addicted to glycolysis and repress their mitochondrial respiration in response to high glucose concentration in their respective media i.e. the Crabtree effect [11, 17].

One method to circumvent this glycolytic addiction and to unmask cancer cells metabolic flexibility *in vitro* is to substitute glucose for galactose. Galactose is metabolised via the Leloir pathway to produce glucose 6-phosphate (G6P), meaning that it can support glycolysis [18]. While glucose is converted into G6P in a one-step reaction by hexokinase (Km ∼ 0.3 mM) [19, 20], galactose requires four biochemical reactions to continuously supply the cell with G6P with the initial reaction catalysed by galactosekinase having a Km value three times higher than hexokinase (galactokinase Km ∼ 1 mM) [21]. This results in galactose providing a slow, yet continuous, supply of carbons, mimicking the TME where glucose is rapidly consumed but continuously being replenished via circulation [22, 23]. Therefore, we used galactose as an *in vitro* tool compound to mimic glycolytic limitation that occurs within the TME.

We have previously shown that cancer cells increase serine catabolism and formate overflow under glycolytic limitation even though biosynthetic demands were decreased [24], indicating a role for one-carbon (1C) cycle in bioenergetics and redox balance [25]. However, the functional advantage and the exact source of carbons for the increased serine catabolism and formate overflow under glycolytic limitation remains unknown. Moreover, it was previously highlighted that glutamine flux is rewired towards serine synthesis pathway (SSP) under glucose starvation via phosphoenolpyruvate carboxykinase 2 (PEPCK2) [26-29]. Both observations indicate that metabolic rewiring takes place towards SSP under glycolytic limitation, yet the exact mechanisms and metabolic advantages of this rewiring remain unclear. Building on these observations, we used stable-isotopic tracers and targeted metabolomics to study the metabolic fluxes of central carbon metabolism and the rewiring that takes place under glycolytic limitation. By using galactose to model glycolytic limitation, we observed a continued but decreased flux through glycolysis while oxygen consumption rates (OCR) were increased [24]. We were able to corroborate previous findings on the rewiring of glutamine metabolism towards SSP via PEPCK2. Surprisingly and despite increased OCR, galactose derived carbons did not enter the TCA cycle for mitochondrial oxidation, suggesting glutamine is the main substrate to support the TCA cycle in these conditions. This block on glycolytic carbons toward mitochondrial metabolism is facilitated by a strong pyruvate kinase isozyme M2 (PKM2) block, pinpointing PKM2 as the rewiring point in glycolysis under glycolytic limitation. Furthermore, we observed that the PKM2 block redirects the limited glycolytic carbons toward SSP to support the survival and proliferation of cancer cells. Finally, we observed an upregulation of the mitochondrial enzymes of 1C cycle where they cooperate to support biosynthesis, bioenergetics and redox balance under glycolytic limitation. These findings reveal that 1C cycle is needed for the proliferation of cancer cells when faced with limited glycolytic supply, and blocking lower glycolysis at PKM2 contributes to keeping the 1C cycle running in these conditions.

## Results

### Cancer cells metabolize galactose and utilize it to support early glycolysis

To ascertain that cancer cells are able to metabolize galactose (Gal), we used a fully labelled [U-^13^C] Gal tracer and compared the relative flux through the Leloir pathway (Fig. 1A) and early glycolysis to the fluxes measured when a [U-^13^C] glucose (Glc) tracer is used (Fig. 1B). The relative flux to UDP-glucose and UDP-galactose was comparable between Glc and Gal tracing and even higher in the case of glucose 1-phosphate in Gal (Fig. 1C-E). When looking at the glycolytic intermediate fructose 1,6-bisphosphate (F 1,6-BP), the ^13^C enrichment was comparable between both tracers suggesting a similar flux of Glc and Gal (Fig. 1F). Pentose phosphate pathway (PPP) is usually highly enriched with ^13^C within the first minutes of [U-^13^C] Glc tracing [30]. Consequently, we saw high enrichments using [U-^13^C] Gal tracing in early PPP intermediate 6-phosphogluconate M+6 (Fig. 1G). However, the ^13^C enrichment in ribose 5-phosphate (R5P) and sedoheptulose 7-phosphate (S7P) showed multiple labelled isotopologues indicating higher shunting of carbons between PPP, early glycolysis and gluconeogenesis in Gal conditions (Fig. 1H, I). Overall, these data demonstrate that cancer cells are able to utilise Gal to fuel early glycolysis and PPP intermediates through the Leloir pathway.

**Figure 1.**
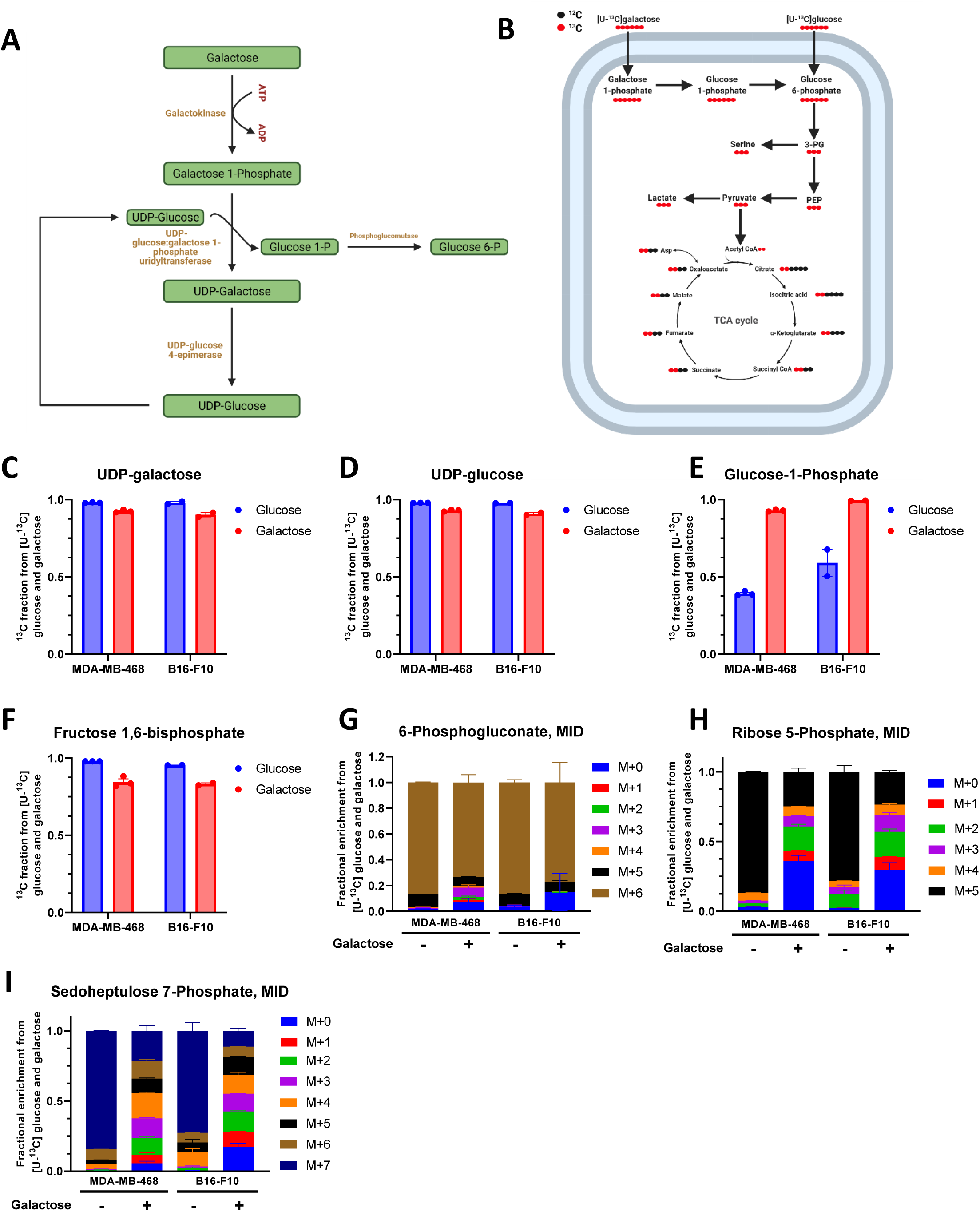
Cancer cells metabolize galactose and utilize it to support early glycolysis. **A** Schematic representation of Leloir pathway for galactose metabolism. **B** Schematic representation of the expected labelling pattern of simplified central carbon metabolism when using [U-^13^C] glucose and [U-^13^C] galactose tracer. **C-F** ^13^C fraction from [U-^13^C] glucose and [U-^13^C] galactose into UDP-galactose **(C)**, UDP-glucose **(D)**, glucose 1-phosphate **(E)** and fructose 1,6-bisphosphate **(F)** after 24h incubation. (**C-F**) Each dot represents an independent experiment averaged from 3 different replicate wells. **G-I** Fractional enrichment in 6-phosphogluconate **(G)**, ribose 5-phosphate **(H)** and sedoheptulose 7-phosphate **(I)** from [U-^13^C] glucose and [U-^13^C] galactose after 24h incubation. Data shown is mean±SEM of 3 (MDA-MB-468) or 2 (B16-F10) independent experiments each being the average of 3 different replicate wells.

### Glutamine and lactate maintain an upregulated oxidative TCA cycle function under glycolytic limitation

To test the validity of using Gal as a tool compound to mimic glycolytic limitation, we sought to first measure the contribution of other carbon sources to the TCA cycle that might support the elevated mitochondrial metabolism under Gal as characterized by increased oxygen consumption rates (OCR) [24] (Fig. S1A). It was previously shown that glutamine anaplerosis into the TCA cycle is increased under full glucose starvation [29, 31]. To test if this is also the case upon limited glycolytic flux (compared to full starvation), we first measured glutamine consumption and observed generally no striking change in the consumption in multiple cancer cell lines under glycolytic limitation (Fig. 2A). Subsequently, we performed [U-^13^C] glutamine tracing in MDA-MB-468, 4T1 and B16-F10 cell lines in Glc and Gal. Upon Gal, we noted an increased relative flux from glutamine into the TCA cycle metabolites in all three cell lines (Fig. 2B-D), indicating that while the absolute consumption was not changed, the usage of glutamine flux is altered. Under Glc, we observed that glutamine enrichment in the TCA cycle metabolites decreased as we move away from glutamine. In contrast, the enrichment remains unchanged under Gal (Fig. 2B-D), indicating that the TCA cycle is running with strongly reduced carbon input from glycolysis. Yet, acetyl-CoA input is required to sustain glutamine oxidation through the TCA cycle, hinting towards malic enzyme (ME) activity to sustain pyruvate levels under glycolytic limitation. ME decarboxylates malate into pyruvate and indeed, we observed a substantial M+3 labelling of pyruvate and lactate from [U-^13^C] glutamine under glycolytic limitation (Fig. 2E and S1B). ME involvement is further supported by an increase in citrate M+6 isotopologue enrichment from glutamine under Gal which suggests increased usage of glutamine derived acetyl-CoA for citrate (Fig. 2F-H). Additionally, when looking at the ratio of M+3 pyruvate/M+4 malate (ME flux), we observe a significant increase under glycolytic limitation (Fig. 2I) [31]. These findings display the metabolic plasticity of cancer cells, as they are able to rewire their metabolism to run the TCA cycle mainly on glutamine. However, we are still able to detect an unlabelled fraction in citrate (Fig. 2F-H) which most likely results from fatty acid oxidation.

**Figure 2.**
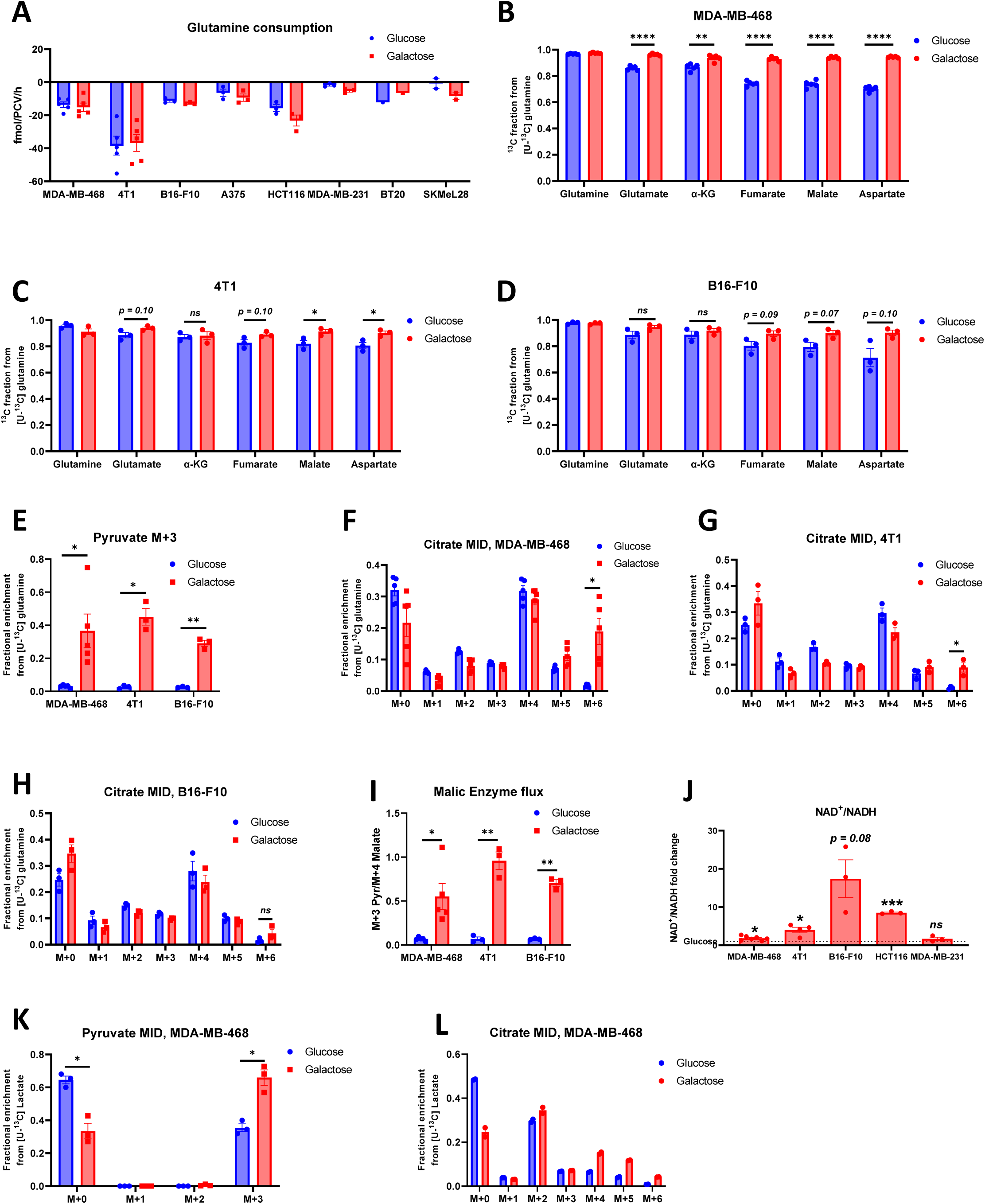
Glutamine and lactate maintain upregulated oxidative TCA cycle function in reliance on malic enzyme (ME) **A** Absolute quantification of glutamine consumption from multiple cancer cell lines after 24h incubation with either glucose or galactose medium. **B-D** ^13^C fraction in TCA cycle metabolites from [U-^13^C] glutamine after 24h incubation with either glucose or galactose medium in MDA-MB-468 **(B)**, 4T1 **(C)** and B16-F10 **(D)**. **E** Fractional enrichment in M+3 pyruvate from [U-^13^C] glutamine after 24h incubation with either glucose or galactose medium in MDA-MB-468, 4T1 and B16-F10. **F-H** Fractional enrichment in citrate from [U-^13^C] glutamine after 24h incubation with either glucose or galactose medium in MDA-MB-468 **(F)**, 4T1 **(G)** and B16-F10 **(H)**. **I** Malic enzyme flux ratio calculated as M+3 pyruvate/M+4 malate isotopologues from [U-^13^C] glutamine after 24h incubation with either glucose or galactose medium in MDA-MB-468, 4T1 and B16-F10. **J** NAD^+^/NADH ratio measured with LC-MS in the indicated cancer cell lines after 24h incubation with galactose and normalised to the same ratio with 24h incubation with glucose medium. **K-L** Fractional enrichment in pyruvate **(K)** and citrate **(L)** from 5mM [U-^13^C] sodium lactate after 24h incubation with either glucose or galactose medium in MDA-MB-468. (**A-L**) Each dot represents an independent experiment averaged from 3 different replicate wells unless stated otherwise. (**A-L**) Mean±SEM, * *p*<0.05, ** *p*<0.01, *** *p*<0.001, **** *p*<0.0001, ns *p*>0.1. *P* value is calculated by Student’s t-test, two-tailed, unpaired with Welch’s correction unless stated otherwise.

An additional carbon source that we might neglect with our *in vitro* setup is lactate whose build up is a prominent feature of most solid tumours [32]. In fact, recent evidence from *in vivo* studies suggest that lactate can be a major carbon source to the TCA cycle of cancer cells [33, 34]. To test if this feature can be recapitulated in our *in vitro* model, we performed [U-^13^C] lactate tracing at a concentration of 5 mM as this was indicated as a physiological relevant concentration within the TME [22, 35]. Supported by the increase in NAD^+^/NADH ratio (Fig. 2J), we measured increased ^13^C contribution to pyruvate and the TCA cycle under glycolytic limitation from [U-^13^C] lactate (Fig. 2K, 2L and S1C, S1D). Of note, the lactate derived carbon contribution to the TCA cycle was mainly via PDH and not via pyruvate carboxylase (Fig S1E, S1F). In summary, we conclude that glutamine and lactate can substitute for glucose to support and run lower glycolysis and the TCA cycle. In the case of lactate, this is facilitated by the increased NAD^+^/NADH ratio favouring lactate oxidation. Other carbon sources might also contribute towards the TCA cycle, however, these contributions are expected to be smaller in comparison to the sources highlighted in this study [33]. Finally, the sustained PDH flux (Fig. S1E) under Gal suggests that a block upstream of PDH prevents glycolysis derived entry into the TCA cycle.

### Glutamine metabolism is redirected towards SSP under glycolytic limitation via PEPCK2

It has previously been shown that under full glucose starvation, gluconeogenesis enzyme PEPCK2 becomes active and supports the biosynthesis of glycolytic intermediates [26-29, 36]. To test if this is also the case in Gal and not only upon full hexose starvation, we monitored glutamine derived carbon contribution to phosphoenolpyruvate (PEP) and 3-PG (Fig. S2A-B) and observed an active PEPCK flux also upon low glycolytic flux. Furthermore, we observed that the ^13^C enrichment was similar under serine/glycine (SG) starvation conditions. Overall, we were able to confirm PEPCK activity in three different cancer cell lines with up to 30% of the measured serine being fully labelled under SG starvation (Fig. 3A). Even with SG in the medium, the glutamine flux to serine under Gal was comparable to the one from Glc (Fig. 3B). As serine is the major substrate for 1C metabolism, we measured the flux from glutamine into vital downstream metabolites of 1C metabolism. We observed ^13^C enrichment in ATP (Fig. 3C), glutathione (GSH) (Fig. 3D) and formate (Fig. S2C). Of note, the labelled fraction of ATP reached an astonishing 40-50% under SG starvation in all three cell lines (Fig. 3C).

**Figure 3.**
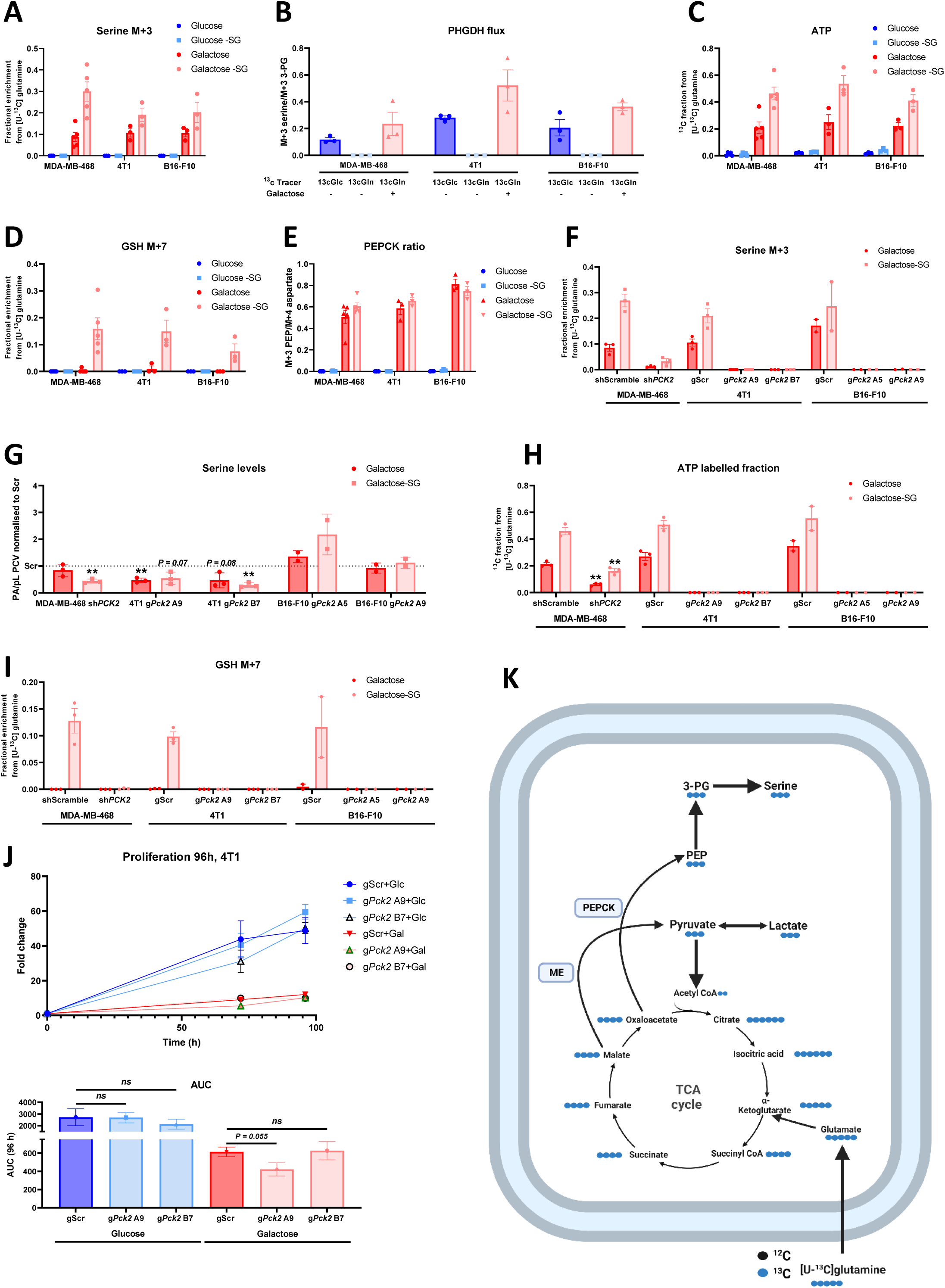
Glutamine metabolism is redirected towards SSP under glycolytic limitation via PEPCK. **A** Fractional enrichment in M+3 serine from [U-^13^C] glutamine after 24h incubation with either glucose ± serine and glycine (SG) or galactose ± SG medium in MDA-MB-468, 4T1 and B16-F10. **B** Phosphoglycerate dehydrogenase (PHGDH) enzyme flux ratio calculated as M+3 serine/M+3 3-PG isotopologues from [U-^13^C] glucose or glutamine (either with glucose or galactose medium) after 24h incubation in MDA-MB-468, 4T1 and B16-F10. **C** ^13^C fraction in ATP from [U-^13^C] glutamine after 24h incubation with either glucose ± SG or galactose ± SG medium in MDA-MB-468, 4T1 and B16-F10. **D** Fractional enrichment in M+7 glutathione (GSH) from [U-^13^C] glutamine after 24h incubation with either glucose ± SG or galactose ± SG medium in MDA-MB-468, 4T1 and B16-F10. **E** Phosphoenolpyruvate carboxykinase (PEPCK) enzyme flux ratio calculated as M+3 PEP/M+4 aspartate isotopologues from [U-^13^C] glutamine after 24h incubation with either glucose ± SG or galactose ± SG medium in MDA-MB-468, 4T1 and B16-F10. **F** Fractional enrichment in M+3 serine from [U-^13^C] glutamine after 24h incubation with either glucose ± SG or galactose ± SG medium in MDA-MB-468 shScramble and sh*PCK2* cells, 4T1 and B16-F10 gScramble and g*Pck2* cells. **G** Peak areas (PA) of intracellular serine in PEPCK2 KD/KOs normalised to packed cell volume (PCV) in picolitres (pL) and presented as fold change to the levels in scramble cells. Data shown is for the levels measured in galactose ± SG medium. **H** ^13^C fraction in ATP from [U-^13^C] glutamine after 24h incubation with galactose ± SG medium and the comparison between scramble and PEPCK2 KD/KOs cells. **I** Fractional enrichment in M+7 glutathione (GSH) from [U-^13^C] glutamine after 24h incubation with galactose ± SG medium and the comparison between scramble and PEPCK2 KD/KOs cells. **J** 2D proliferation curve of 4T1 gScramble and g*Pck2* cells with either glucose or galactose, with the corresponding area under the curve (AUC). 72h timepoint includes 3 independent experiments while 96h includes 4 independent experiments (from at least 3 different replicate wells for each independent experiment). (**A-J**) Each dot represents an independent experiment averaged from 3 different replicate wells unless stated otherwise. (**A-J**) Mean±SEM, * *p*<0.05, ** *p*<0.01, *** *p*<0.001, **** *p*<0.0001, ns *p*>0.1. *P* value is calculated by Student’s t-test, two-tailed, unpaired with Welch’s correction unless stated otherwise. **K** Schematic representation of the rewiring of glutamine metabolism that take place under glycolytic limitation.

PEPCK catalyses the decarboxylation of oxaloacetate into PEP, therefore, we can measure its metabolic flux by using aspartate as a surrogate for oxaloacetate as the tracer medium contains no aspartate [37]. Doing so, we observed a clear increase in PEPCK flux under glycolytic limitation with and without SG (Fig. 3E). Mitochondrial PEPCK (PEPCK2) has been identified as the major isoform present in most tumour tissues [26]. Here we observed that PEPCK2 protein expression was markedly increased under glycolytic limitation in all the cell lines tested (Fig. S2D). Subsequently, we used two different genetic approaches to target PEPCK2; we knocked down PEPCK2 in MDA-MB-468 cells using shRNA and knocked it out using CRISPR/Cas9 system in 4T1 and B16-F10 cells (Fig. S2E, F). Overall, we were able to confirm PEPCK2 as the culprit for the ^13^C enrichment into serine from glutamine (Fig. 3F). More importantly, serine intracellular levels significantly dropped in MDA-MB-468 and 4T1 but not in B16-F10, signifying the importance of the glutamine flux for the serine pool while also highlighting varying cell line specific dependencies on PEPCK2 (Fig. 3G). Furthermore, the labelling into ATP and GSH was significantly reduced in PEPCK2 KD and eliminated in the KOs (Fig. 3H, I), while formate overflow was unchanged in the PEPCK2 KOs (Fig. S2G). Moreover, we observed a mild increase in the labelling of TCA cycle intermediates and pyruvate from glutamine in PEPCK2 KD/KOs which most likely is a result of the elimination of the flux through PEPCK (Fig. S2H, I). Despite the significance of the glutamine flux into SSP, we noted no difference in proliferation of 4T1 PEPCK2KO under glycolytic limitation (Fig. 3J). This indicates that 4T1 cells are able to compensate for the missing contribution into serine by either increasing uptake of serine from the medium or increasing the flux from Gal under SG starvation (Fig. S2J). Overall, by using Gal as a tool compound, we were able to see that glutamine metabolism is rewired in cancer cells to run the TCA cycle independently and to support SSP via enhanced activity of PEPCK2 (Fig. 3K). However, it remains unclear how PEPCK2 and ME fluxes are coordinated as these fluxes cannot be disentangled by using [U-^13^C] glutamine. Therefore, we opted to use [U-^13^C] Gal as a tracer to clearly follow the glycolytic flux into lower glycolysis and TCA cycle under glycolytic limitation.

### Cancer cells sustain carbon flux towards SSP under glycolytic limitation

To study the relative fluxes of lower glycolysis under glycolytic limitation in more detail we used [U-^13^C] Gal. Across a panel of nine different cancer cell lines representing four different tumours (breast, colorectal, glioblastoma and melanoma) we observed robust Gal metabolization through glycolysis and SSP (Fig. 4A-C). Despite similar enrichment in F 1,6-BP (Fig. 1F), the relative flux of ^13^C in Gal decreased towards late glycolytic intermediates with especially pyruvate showing only about 10% of the ^13^C enrichment observed under Glc, which could be explained partly by the increased flux from [U-^13^C] glutamine under Gal (Fig. 4B). In comparison, cells appeared to keep about 25 – 50% of their relative flux towards serine under Gal (Fig. 4C). To exclude that this effect is due to differences in the time needed to reach isotopic steady state [37], we performed a time course experiment. We observed isotopic steady state conditions with both tracers at around 24h and longer incubation times did not result in higher ^13^C enrichment of pyruvate, suggesting that the different labelling pattern in pyruvate is not a result of different flux rates but of an alteration of central carbon metabolism upon Gal (Fig. S3A). Furthermore, different Gal concentrations also did not affect how much [U-^13^C] Gal enriched the pyruvate pool (Fig. S3A) nor did the usage of RPMI-1640 (Fig. S3B). Using dialysed or normal FBS did not affect pyruvate enrichment as well (Fig. S3C). To test the [U-^13^C] Gal enrichment in a more physiological setting, we repeated the ^13^C tracing in Plasmax medium [38] and observed similar reduction in M+3 pyruvate isotopologue between Gal and Glc (Fig. S3D). To eliminate the possibility of an inhibitory action of Gal, we mixed Glc and Gal in a 1:1 mix where the Glc added was ^13^C. If Gal had a direct inhibitory action on glycolysis we would expect to see a clear reduction of ^13^C flux towards pyruvate, however, we observed no difference in ^13^C enrichment in glycolysis under such conditions (Fig. S3E). These observations indicate that the reduced glycolytic ^13^C flux is a result of the cells’ response to the limited availability of carbons under Gal.

**Figure 4.**
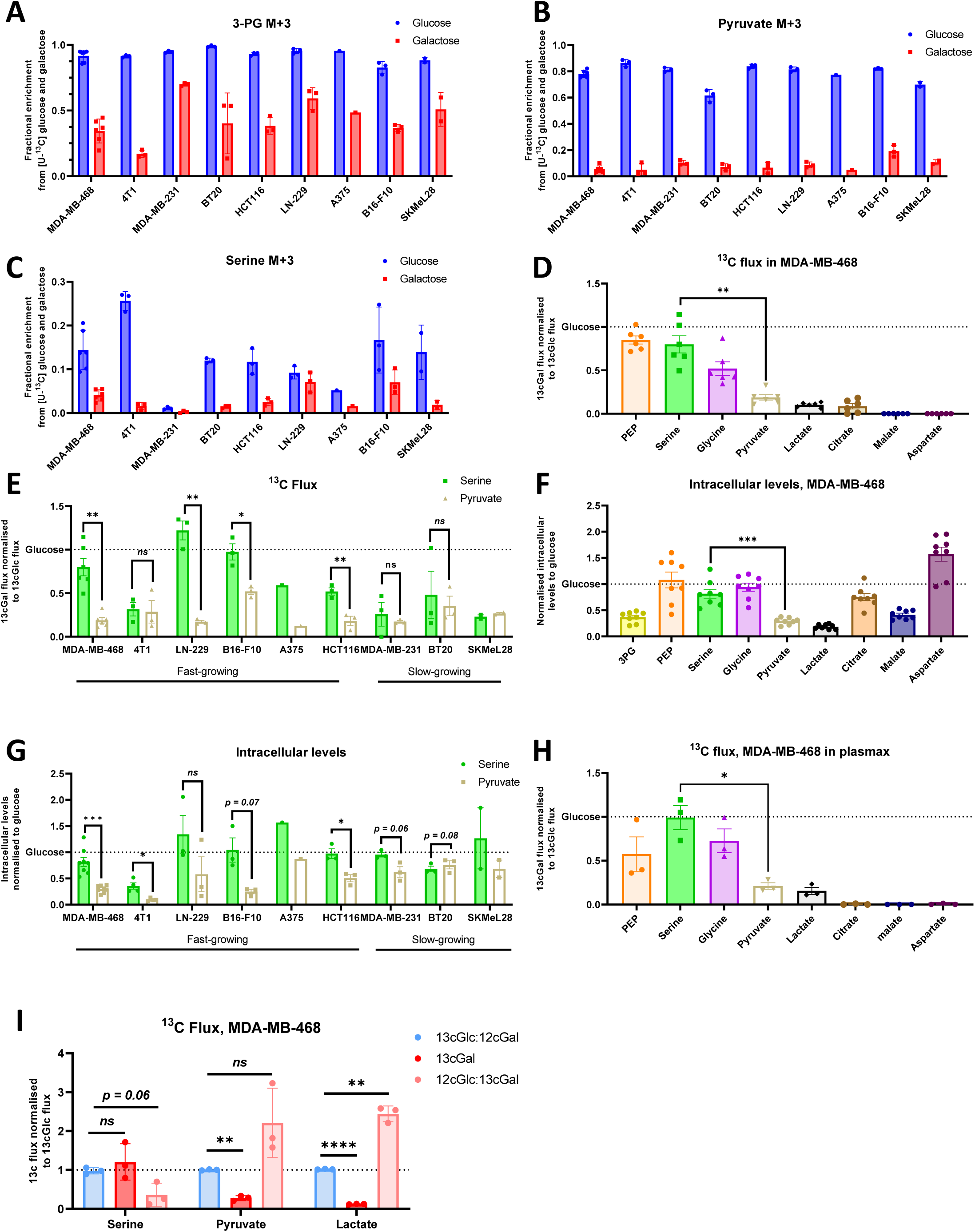
Cancer cells sustain carbon flux towards serine synthesis pathway (SSP) under glycolytic limitation. **A-C** Fractional enrichment from 17.5 mM [U-^13^C] glucose and galactose into M+3 3PG (**A**), M+3 pyruvate (**B**) and M+3 serine (**C**) after 24h incubation measured with LC-MS. **D-E** The normalised relative ^13^C flux under galactose towards multiple metabolites represented as a fold change of the same ratio under glucose in MDA-MB-468 (**D**) and a panel of 9 cancer cell lines (**E**). Fast-growing cells are those with doubling time <24h. **F-G** Intracellular levels of multiple metabolites under galactose represented as a fold change of the levels under glucose in MDA-MB-468 (**F**) and a panel of 9 cancer cell lines (**G**). **H** The normalised relative ^13^C flux under galactose towards multiple metabolites represented as a fold change of the same ratio under glucose in MDA-MB-468 when cultured in physiological medium Plasmax. **I** The normalised relative ^13^C flux under galactose and glucose:galactose mixes towards multiple metabolites represented as a fold change of the same ratio under glucose in MDA-MB-468. (**A**-**I**) Each dot represent an independent experiment (from 3 different replicate wells). (**A-I**) Mean±SEM, * *p*<0.05, ** *p*<0.01, *** *p*<0.001, **** *p*<0.0001, ns *p*>0.1. *P* value is calculated by Student’s t-test, two-tailed, unpaired with Welch’s correction.

To better visualise the differences between the relative flux towards lower glycolysis and SSP, we used 3-PG (divergent point between SSP and lower glycolysis) M+3 isotopologue as a common denominator to normalise the main labelled isotopologue of different metabolites (e.g. M+3 serine/M+3 3PG, M+3 pyruvate/M+3 3PG, M+2 citrate/M+3 3PG etc). The resulting ratios are presented as a fold change of the same ratios under Glc (Fig. 4D, E). We found that in most cell lines the ratio for serine under Gal was comparable or higher than under Glc, which indicates comparable or higher flux to SSP under glycolytic limitation. In contrast, the ratio of M+3 pyruvate / M+3 3PG was strikingly lower under Gal than under Glc, which indicates significantly less carbon flux towards pyruvate under Gal. Thus, we observed that cancer cells re-wire their limited ^13^C supply under Gal towards SSP resulting in SSP as the significantly favoured destination compared to lower glycolysis. This phenotype was predominant in cell lines which doubled under 24h (fast-growing) (Fig. 4D, E). One exception was the murine breast cancer cell line 4T1, which is extremely glycolytic and is very sensitive to Gal, indicating that this cell line may not be able to adapt to such an environment (Fig. 4E). Furthermore, we took note of a similar observation of the rewiring when looking at the intracellular levels of serine and pyruvate (Fig. 4F, G). Subsequently, we validated the re-wiring phenotype when MDA-MB-468 were cultured in the physiological medium Plasmax (Fig. 4H). When ^13^C-Glc was mixed 1:1 with ^12^C-Gal, we saw that there was no change in the ^13^C flux ratios, showing again that Gal is not inhibitory of glycolysis (Fig. 4I). However, when ^13^C-Gal was mixed with ^12^C-Glc we saw that the flux was heavily favoured towards pyruvate and lactate and not to SSP (Fig. 4I). This indicates that under ample supply of glycolytic carbons, ^13^C flux from Gal is not rewired towards SSP.

Based on the observed rewiring, we checked the expression of SSP enzymes and observed a modest increase in the protein expression of phosphoserine aminotransferase 1 (PSAT1) and phosphoserine phosphatase (PSPH), while phosphoglycerate dehydrogenase (PHGDH) expression was mostly not elevated (Fig. S3F, S3G). This indicates that the sustained flux through SSP is not a result of altered gene expression but rather a more direct metabolic adaptation of enzyme activity. In summary, under glycolytic limitation, fast-growing cancer cells (cell lines with doubling times <24 hours) re-wire their metabolism upon limited glycolytic flux towards SSP at the cost of pyruvate oxidation.

### Glycolytic limitation inhibits glycolysis at PKM2

Having identified a block at lower glycolysis and re-routing of glycolytic carbons towards SSP, we looked to determine how cancer cells mechanistically undergo the proposed re-wiring. Since the re-wiring occurs somewhere between 3-PG and pyruvate, we hypothesised that this might occur at pyruvate kinase which is a known regulatory node of glycolysis [39] (Fig. 5A). This hypothesis is supported by the observation that under Gal cancer cells excrete almost no lactate (Fig. S4A) and its intracellular levels drop significantly (Fig. 4F). This finding also explains why cancer cells are very vulnerable to adenine nucleotide translocator (ANT) inhibition under glucose starvation [40] as in the absence of lactate production they would be highly relying on mitochondrial ATP. Specifically, we have observed that PEP accumulates under Gal while pyruvate levels drop dramatically (Fig. 5B-E), further supporting the hypothesis that the metabolic block occurs at the level of pyruvate kinase. This is further substantiated by the observation that ^13^C labelling in pyruvate is nearly absent across all cell lines analyzed, while PEP remains significantly labelled (Fig. 5B-E). To confirm the glycolytic block at PKM2, we used the PKM2 activator TEPP-46 [41] and repeated stable isotope tracing with [U-^13^C] Gal. The activation of PKM2 increased the ^13^C flux towards citrate at the cost of serine synthesis (Fig. 5F). Furthermore, we observed a significant reduction of the labelling into serine from [U-^13^C] glutamine solidifying our model that the rewiring into SSP is supported by PKM2 inhibition (Fig. S4B). Finally, the activation of PKM2 and subsequent prevention of the rewiring towards SSP lead to a significant decrease in proliferation of MDA-MB-468 (Fig. 5G) and reduction in 3D growth using anchorage-independent conditions (Fig. S4C), concluding that maybe not the PEPCK2 flux but the inhibition of the PKM2 flux is important for survival, at least in our experimental setup (Fig. 3J, 5G). F 1,6-BP binds to PKM2 and induces the formation of the active tetramer form [42, 43]. Under Gal we saw that the intracellular levels of F 1,6-BP drop (Fig. S4D) which is known to lead to a reduction in PKM2 activity [42, 43]. However, what we observe with Gal is a complete block of the glycolytic flux at PKM2, indicating that factors other than the low levels of F 1,6-BP are contributing to this rewiring.

**Figure 5.**
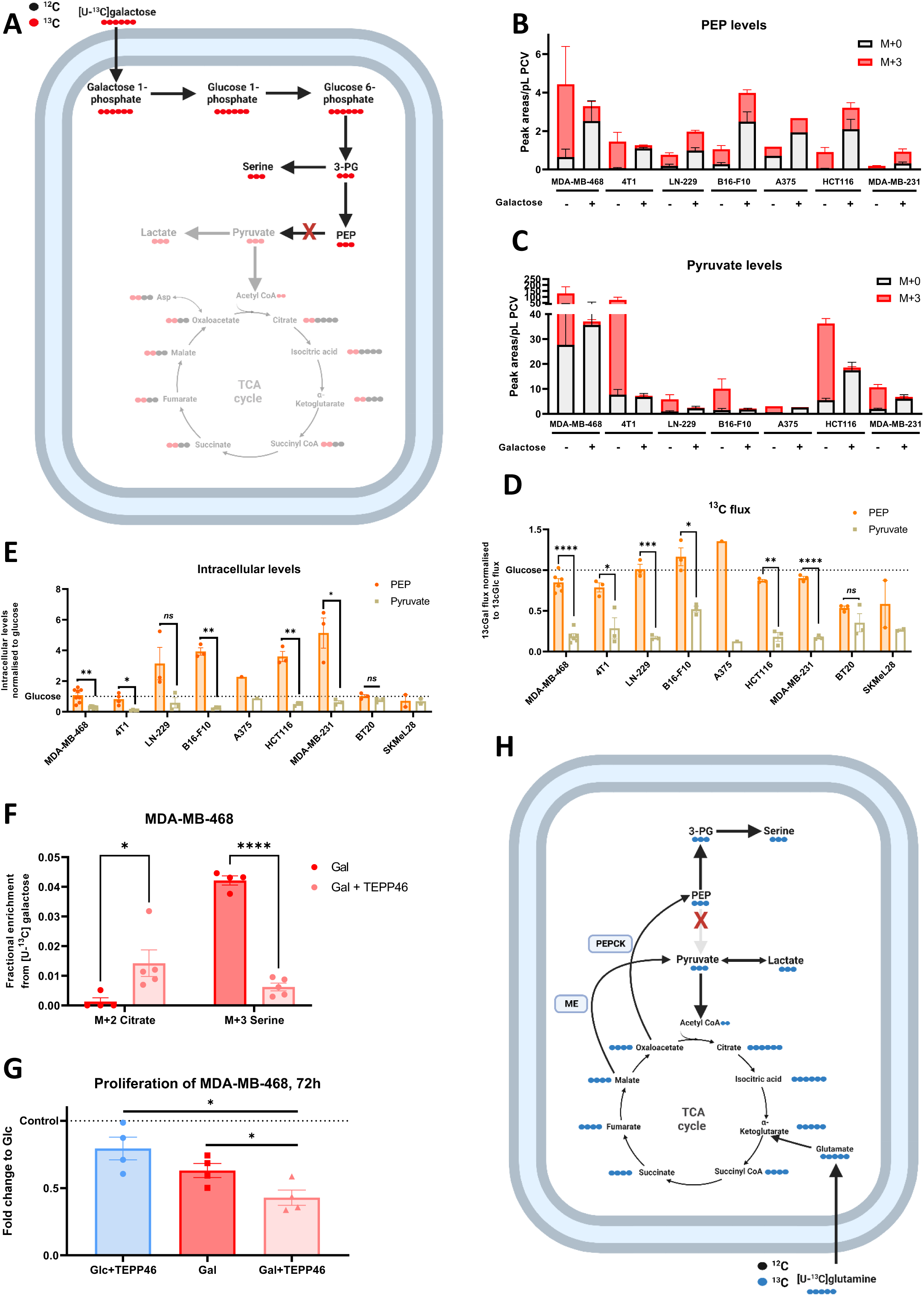
Glycolytic limitation inhibits glycolytic flux at pyruvate kinase isozyme M2 (PKM2) **A** Schematic representation of glycolysis and TCA cycle upon isotopologue labelling with [U-^13^C] glucose and galactose with the proposed block at PKM2. **B-C** Peak areas of M+0 and M+3 isotopologues of PEP (**B**) and pyruvate (**C**) after 24h incubation with [U-^13^C] glucose and galactose and normalisation to packed cell volume (PCV). **D** The normalised relative ^13^C flux under galactose towards PEP and pyruvate represented as a fold change of the same ratio under glucose in a panel of 9 cancer cell lines. **E** Intracellular levels of PEP and pyruvate under galactose represented as a fold change of the levels under glucose in a panel of 9 cancer cell lines. **F** Fractional enrichment from [U-^13^C] galactose into M+2 citrate and M+3 serine after 24h incubation (Gal+TEPP-46) as measured with LC-MS. **G** 2D proliferation of MDA-MB-468 treated with 100 µM of TEPP-46 with either glucose or galactose and measured as cell counts at 72h relative to proliferation under glucose. (**B**-**G**) Each dot represents an independent experiment (from at least 3 different replicate wells). (**B**-**G**) Mean±SEM, * *p*<0.05, ** *p*<0.01, *** *p*<0.001, **** *p*<0.0001, ns *p*>0.1. *P* value is calculated by Student’s t-test, two-tailed, unpaired with Welch’s correction. **H** Schematic representation of the rewiring of glutamine metabolism that take place under glycolytic limitation and the differentiation between ME and PEPCK2 fluxes.

In summary, we provide so far evidence that when hexose availability for glycolysis is limited, the residual glycolytic carbon remains in upper glycolysis to support SSP. Meanwhile, glutamine and lactate compensate for carbon demand of the TCA cycle to support increased mitochondrial respiration. Moreover, we are able to increase the mechanistic granularity of the glutamine rewiring through ME and PEPCK by identifying the PKM2 block which means that ME flux (Gln --> Pyruvate) is exclusive to supporting the TCA cycle while PEPCK flux (Gln --> PEP) is exclusive to support SSP (Fig. 5H).

### SSP is needed for proliferation and viability under glycolytic limitation

Next, we sought to investigate the functional advantage of the observed metabolic re-wiring towards SSP. We noted a weak correlation between the growth rate of cancer cells under Glc and to what extent Gal inhibited their proliferation (Fig. 6A). Based on that observation we studied the proliferative capacity of the MDA-MB-468 cell line upon treatment with a competitive inhibitor of PHGDH (the rate-limiting enzyme of SSP) BI-4916 [44] while under glycolytic limitation. We observed the lowest proliferative capacity of MDA-MB-468 cells when SSP was inhibited under glycolytic limitation (Fig. 6B). MDA-MB-468 cells grown in 3D using anchorage independent conditions with both normal and dialysed FBS also showed the significant growth reduction upon SSP inhibition and glycolytic limitation (Fig. 6C, D and S5A). We further confirmed these results in different cancer cell lines with similar results for two out of three cell lines tested (Fig. 6E). Lastly, we also analyzed the viability of different cancer cell lines in Gal treated with BI-4916 and compared the fold-change apoptosis induction to Glc cultivation in different cell lines. We found an increased percentage of apoptotic (Ann V^+^) cells when PHGDH flux was blocked under glycolytic limitation (Fig. 6F). The exception is here again the 4T1 cell line, a finding that matches the fact this was the only cell line not showcasing the observed rewiring towards SSP under glycolytic limitation (Fig. 4E). Overall, it is evident that sustained SSP exerts a pro-proliferative and anti-apoptotic role under glycolytic limitation.

**Figure 6.**
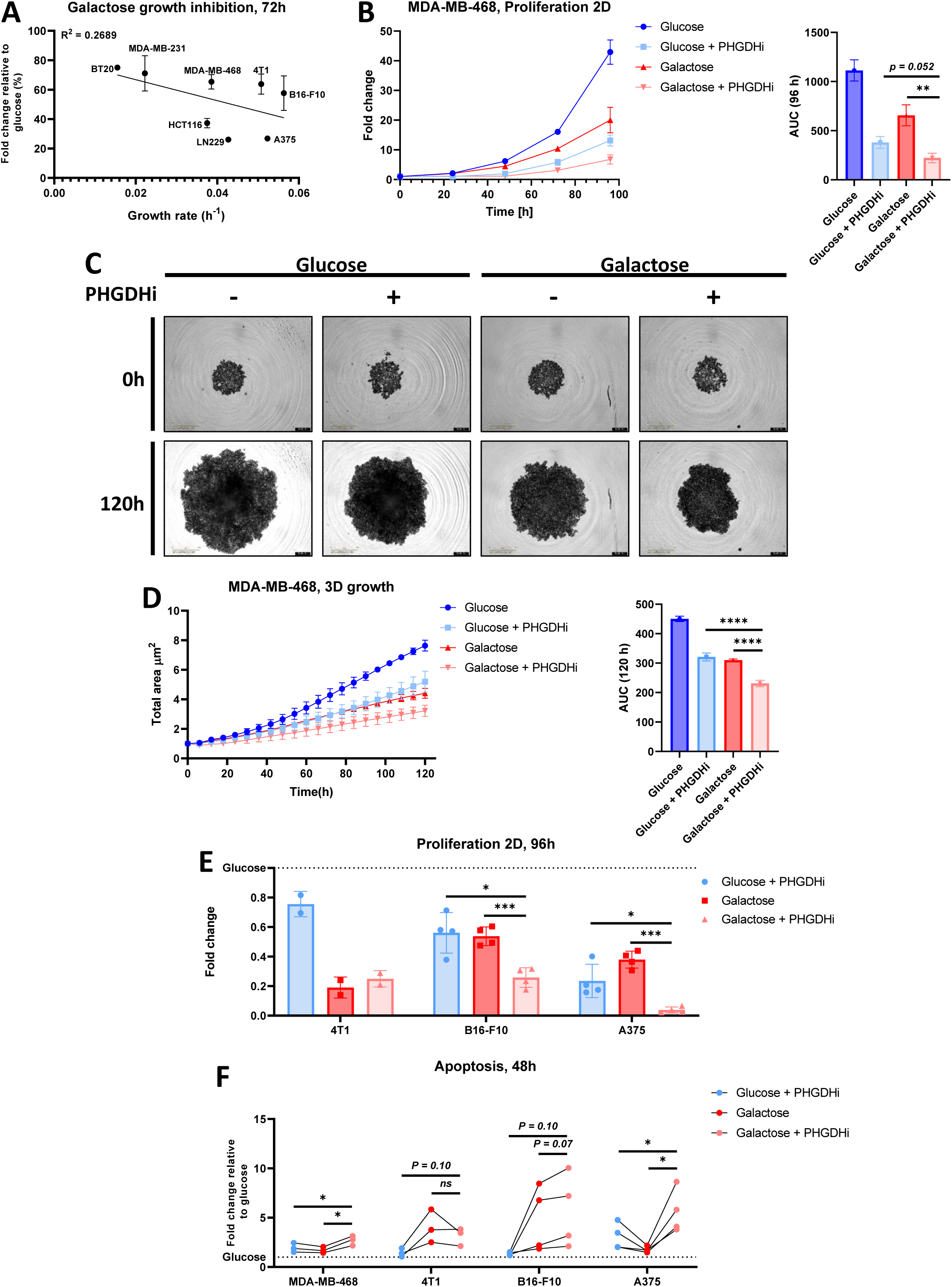
SSP is needed for proliferation and viability under glycolytic limitation. **A** The correlation between the growth rate of different cancer cell lines and the extent to which galactose inhibits their proliferation (represented as a percentage of growth under glucose). MDA-MB-468 n=5, 4T1 n=2, HCT116 n=2, MDA-MB-231 n=2, B16-F10 n=3, A375 n=3, BT20 n=1 and LN-229 n=1. R^2^ measured for the line of best-fit using simple linear regression. **B** 2D proliferation of MDA-MB-468 when treated with 15 µM of the PHGDH inhibitor BI-4916 (PHGDHi) with either glucose or galactose, with the corresponding area under the curve (AUC). 24h and 96h timepoints include 4 independent replicates while 48h and 72h include 5 independent replicates (from at least 3 different replicate wells for each independent experiment). **C** Bright field images are representative of 3 independent experiments of MDA-MB-468 aggregates growing in anchorage-independent conditions. Scale bar represent 800 µm. **D** 3D growth of MDA-MB-468 aggregates using anchorage-independent conditions when treated with 15 µM of PHGDHi with either glucose or galactose, with the corresponding AUC. 3 independent experiments are represented each with at least 6 different replicate wells. **E** 2D proliferation of multiple cancer cell lines when treated with 15 µM of PHGDHi with either glucose or galactose as measured with cell counts at 96h relative to proliferation under glucose. Each dot represent an independent experiment from at least 3 different replicate wells. (**B**, **D, E**) Mean±SEM, * *p*<0.05, ** *p*<0.01, *** *p*<0.001, **** *p*<0.0001, n.s. *p*>0.1. *P* value is calculated by Student’s t-test, two-tailed, unpaired with Welch’s correction. **F** Viability of different cancer cell lines when treated with 15 µM of PHGDHi with either glucose or galactose for 48h as measured as annexin V^+^ cells after excluding necrotic cells (PI). MDA-MB-468 and 4T1 n=3, B16-F10 and A375 n=4. Data shown is the mean, * *p*<0.05, ns *p*>0.1. *P* value is calculated by Student’s t-test, two-tailed, paired.

### SSP flux is crucial to sustain mitochondrial formate production and to support all three metabolic pillars

In addition to serine synthesis, we were interested to understand the functional relevance of sustained serine availability. We sought to uncover the metabolic purpose of such rewiring, especially since 1C metabolism is able to support all of the three metabolic pillars; biosynthesis, bioenergetics and redox balance [25] (Fig. 7A). We analyzed the expression of the different enzymes of the 1C cycle and found that the mitochondrial enzymes SHMT2 and MTHFD2 displayed increased mRNA expression under glycolytic limitation while the expression of their cytoplasmic versions were mostly kept unchanged with MTHFD2 expression also increased at the protein level (Fig. 7B-C). The product of the mitochondrial 1C cycle branch is the release of the 1C unit as formate and we have previously shown that serine catabolism and formate overflow increase upon Gal [24]. In corroboration, we performed formate measurements in a panel of cancer cell lines and observed increased formate release under glycolytic limitation across the cell lines tested (Fig. 7D). Similar to lactate overflow, formate overflow produces one ATP molecule per MTHFD1L reaction. In addition to ATP production, formate overflow yields one NADH molecule via MTHFD2 [45]. The coupling of MTHFD2-derived NADH to the electron transport chain (ETC) thereby contribute to mitochondrial ATP production and the observed increase in OCR upon Gal (Fig. S1A)[24]. Meaning that the increased formate release can serve as a crucial bioenergetic function by supporting mitochondrial ATP production under glycolytic limitation [24]. Furthermore, we have previously shown that under glycolytic limitation, AMPK is activated alongside the increased formate release [24] which is an observation we could confirm in the glioblastoma cell line LN-229 (Fig. S6A). To test whether AMPK is responsible for the rewiring towards SSP and 1C cycle, we utilized LN-229 cells harbouring a genetic deletion of both alpha subunits of AMPK (Fig. S6B) [46]. We observed no difference in the AMPKKO cells in terms of formate release (Fig. S6C) and ^13^C flux rewiring (Fig. S6D), indicating that AMPK activation and increased formate overflow are not in causal relation. Furthermore, knocking-out MTHFD1L, the enzyme catalysing formate release, led to only a subtle decrease in growth of MDA-MB-468 cells in 2D and 3D anchorage-independent conditions under glycolytic limitation (Fig. S6E-G).

**Figure 7.**
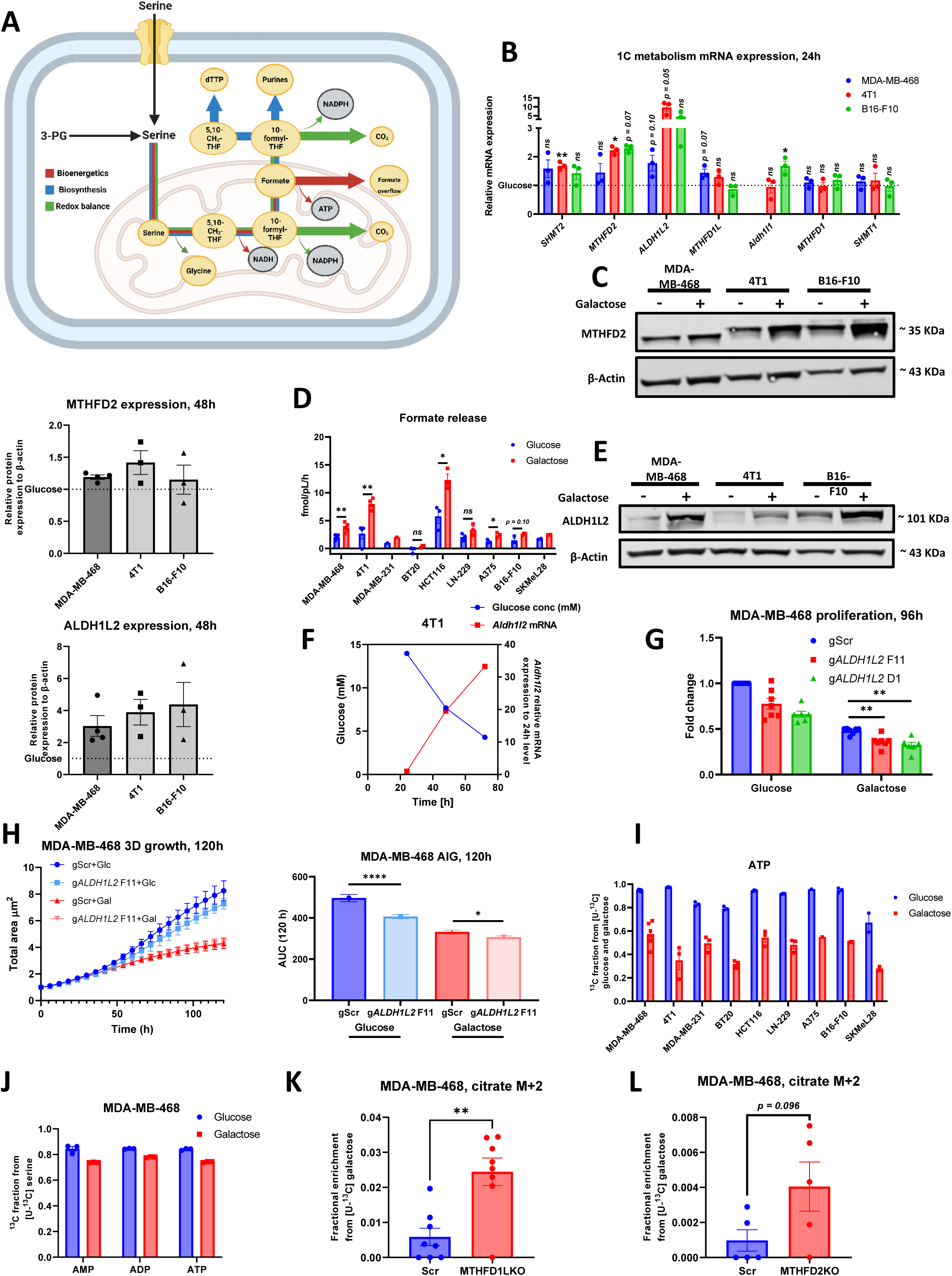
SSP flux is crucial to sustain mitochondrial formate production to support all three metabolic pillars. **A** Schematic representation of the one-carbon cycle with the different pillars and how it can contribute to biosynthesis, bioenergetic and redox balance. **B** mRNA expression of *SHMT2*, *MTHFD2*, *AALDH1L2*, *MTHFD1L*, *Aldh1l1, MTHFD1* and *SHMT1* genes relative to *GAPDH* and *ACTINβ* or *Gapdh and Sdha* after 24h incubation with either glucose or galactose measured with RT-qPCR. Data presented as fold change to the mRNA expression in cells cultured with glucose. **C** Protein expression of MTHFD2 after 48h incubation with either glucose or galactose, with the quantification of these signals relative to β-actin as a loading control and presented as a fold change of expression under galactose to that of glucose. **D** Absolute quantification of formate release from the indicated cancer cell lines after 24h incubation with either glucose or galactose. **E** Protein expression of ALDH1L2 after 48h incubation with either glucose or galactose, with the quantification of these signals relative to β-actin as a loading control and presented as a fold change of expression under galactose to that of glucose. **F** Concentration of glucose as measured in media in which 4T1 cells were cultured for 72h, compared to A*ldh1l2* mRNA expression relative to *Gapdh* and presented as fold change for expression at 24h. Data presented in **F** is from one independent experiment, mRNA was extracted from single wells while medium measurement are mean±SEM of 3 different replicate wells. **G** 2D proliferation of MDA-MB-468 Scramble and ALDH1L2KO cells cultured in either glucose or galactose as measured with cell counts at 96h relative to proliferation of scramble cells under glucose. **H** 3D growth of MDA-MB-468 control and ALDH1L2KO aggregates using anchorage-independent conditions in either glucose or galactose, with the corresponding AUC. 3 independent experiments are represented each with at least 6 different replicate wells. **I** ^13^C fraction in ATP from [U-^13^C] glucose and galactose after 24h incubation in multiple cell lines. **J** ^13^C fraction in AMP/ADP/ATP from [U-^13^C] serine after 24h incubation in MDA-MB-468. **K-L** Fractional enrichment from [U-^13^C] galactose into M+2 citrate after 24h incubation in MDA-MB-468 Scramble compared to MTHFD1LKO (**K**) and MTHFD2KO (**L**) as measured with LC-MS. (**B-L**) Each dot represent an independent experiment averaged from 3 different replicate wells unless stated otherwise. (**B**-**L**) Mean±SEM, * *p*<0.05, ** *p*<0.01, *** *p*<0.001, **** *p*<0.0001, ns *p*>0.1. *P* value is calculated by Student’s t-test, two-tailed, unpaired with Welch’s correction unless stated otherwise.

Having observed that the bioenergetic aspect of ATP production upon formate overflow might only partially contribute to the functional implications during glycolytic restriction, we investigated potential additional metabolic aspects of the 1C cycle that support redox balance and biosynthesis. The 1C unit donated by serine can either be used for de novo purine and pyrimidine synthesis or can be oxidised to CO_2_ via ALDH1L2 or ALDH1L1 while generating NADPH in the latter reaction (Fig. 7A). NADPH provides the cell with reducing equivalents required to counter reactive oxygen species (ROS). We explored both biosynthesis and redox balance as potential sinks for the rewired ^13^C flux. ALDH1L2 expression was markedly increased under glycolytic limitations in different cancer cell lines while the expression of ALDH1L1 was unchanged at mRNA level (Fig. 7B, 7E). This indicates that 1C metabolism is mostly involved in maintaining mitochondrial rather than cytosolic redox state under glycolytic limitation. Moreover, when we cultivated the highly glycolytic breast cancer cell line 4T1 (Fig. S4A) for 72h in Glc without medium change, we saw that the expression of ALDH1L2 increased as Glc concentration became limited supporting the ALDH1L2 data we obtained using Gal (Fig. 7F). Knocking out ALDH1L2 in MDA-MB-468 cells (Fig. S6H) resulted in mildly reduced proliferation under glycolytic limitation in both 2D and 3D assays (Fig. 7G, H). Concurrent increased expression of ALDH1L2 and increased formate overflow would together highlight the extent of increased carbon flux through SSP and 1C cycle as despite the release of 1C as CO_2_, formate overflow is still significantly increased.

PPP is a key regulator of biosynthesis and redox balance as it supplies the cells with R5P and NADPH [47]. The cycling of carbons we observed (Fig. 1H, I) suggests that PPP is shifted towards maximising the production of NADPH and R5P from the limited carbon supply under Gal. Due to this carbon cycling in PPP and subsequent labelling of R5P, it is not possible to determine the contribution of 1C metabolism to nucleotide biosynthesis by using [U-^13^C] Glc and Gal tracers (Fig 7I and S6I-M). Therefore, we utilised [U-^13^C] serine tracer and observed that the flux from serine towards nucleotide biosynthesis is sustained under glycolytic limitation (Fig. 7J). Overall, it can be concluded that the metabolic functions of the 1C metabolism are individually only partially contributing to the functionality of cells under glycolytic limitation. Conclusively, this suggests the idea that the flux through the mitochondrial 1C cycle might serve multiple metabolic purposes.

To test if mitochondrial 1C metabolism plays a combined role in controlling metabolic adaptation to glycolytic limitation, we performed [U-^13^C] Gal tracing in MTHFD1L and MTHFD2 KO cells (Fig. S6G) and found that the we can detect an increase in labelling into TCA cycle upon loss of mitochondrial 1C cycle activity (Fig. 7K, L). This can be seen in the decrease in ^13^C flux to serine and glycine with concomitant increase of ^13^C flux to citrate (Fig. S6O, P). This effect was clearest in MTHFD1LKO and was similar to what we observed with PKM2 activation (Fig. 5F). These observations indicate that the mitochondrial 1C pathway contribute to the rewiring of lower glycolysis.

Put together, it is apparent that the rewiring towards SSP and the plasticity of 1C metabolism under glycolytic limitation serves all three metabolic pillars with sustained carbon flux from serine to purines and increased ATP and NADPH generation. In particular, mitochondrial formate production needs to be increased, as an increased fraction of formate is likely to be oxidized due to increased protein levels of ALDH1L2 while at the same time formate overflow is also increased. Additionally, cytosolic formate is needed to support purine and pyrimidine biosynthesis to sustain proliferation. Finally, we observed that loss of mitochondrial 1C pathway activity results in ^13^C carbon re-entry through PKM2 towards the TCA cycle.

### *In vivo* relevance of Gal to model glycolytic limitation

The *in vitro* modelling of the metabolic environment that cancer cells face *in vivo* is extremely challenging due to the heterogeneity of the tumour tissue and the varying metabolite composition of the TME. Using Gal as a tool, we aimed to mimic glucose limitation as one important variable of the *in vivo* TME. To test the physiological relevance of using Gal as a tool compound to study glycolytic limitation, we subcutaneously injected B16-F10 cells into the flanks of C57BL/6 mice. We then allowed the tumours to grow for 16 and 9 days, thereby generating larger and smaller tumours, respectively, (Fig. 8A, B) and then we infused the mice under anaesthesia with [U-^13^C] Glc tracer for 2 hours. Next, we performed targeted metabolomics on pieces of dissected tumour and compared the small tumours to the inside and the outside of the big tumours to measure the gradient of metabolite levels between the outer layer and the core of the relatively big tumours. We observed that glucose levels strikingly drop in the core of the big tumour compared to its outer layer (Fig. 8C). Meanwhile, other metabolites that can act as secondary carbon sources for tumours (lactate, glutamine and serine) were all present at similar levels between core and the outer layer (Fig. 8C). Moreover, when looking at the MIDs of 3-PG we can see that the ^13^C enrichment is slightly higher on the outer layer than in the core (Fig. 8D). In tumours, when the ^13^C fraction of pyruvate is normalised to the ^13^C fraction of 3-PG, the ratio is equal to one when newly synthesised pyruvate is entirely generated by glycolysis [34]. The ratio is in contrast >1 when other ^13^C sources outside of glycolysis, that are labelled elsewhere by [U-^13^C] Glc and then supply the tumour from circulation, contribute to the pyruvate pool [34]. Comparing these ratios between the outer and inner layers of the B16-F10 tumours, we can see that the ratio is ∼ 2.3 in the inner core and 1.5 in the outer layer (Fig. 8E). This indicates that the core is depleted of glucose in bigger tumours and cells within this core are relying more on other carbon sources due to altering their glycolytic usage i.e. metabolic flexibility. In contrast, serine ratios remain rather constant suggesting a switch of the glycolytic flux towards serine (Fig. 8E). Even though these data are not definitive proof, they hint towards similar rewiring as observed *in vitro* when culturing cells in Gal.

**Figure 8.**
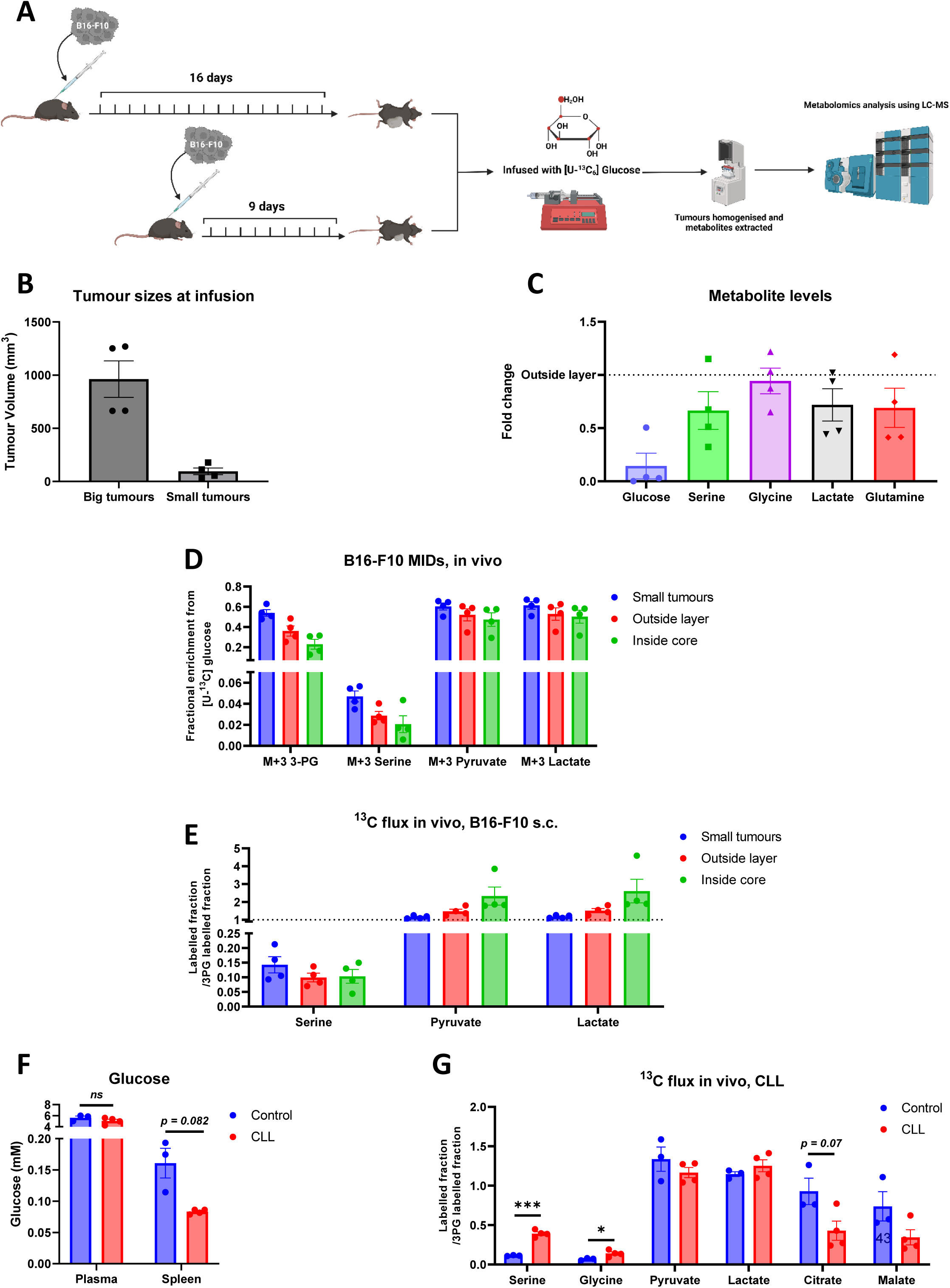
*In vivo* relevance of using Gal to model glycolytic limitation. **A** Schematic representation of the *in vivo* setup for B16-F10 subcutaneous tumours. **B** Tumour volume of B16-F10 tumours grown subcutaneously on the flanks of C57BL/6 mice (n=4/group). **C** Metabolite levels measured from the inside core of B16-F10 tumours and presented as fold change of the level of those metabolites in the outside layer of the same tumours. **D** Fractional enrichment in tumour piece of the presented metabolites after 2h infusion with [U-^13^C] glucose via the tail vain. **E** The ^13^C flux from 3-PG towards multiple metabolites measured in B16-F10 tumours. **F** Glucose concentration as measured in the plasma and in grinded spleen tissue from control and CLL mice after 2h continuous infusion with [U-^13^C] glucose. The concentration was determined using standard addition method. **G** The ^13^C flux from 3-PG towards multiple metabolites measured in CLL cells collected from the spleen of CLL mice. (**B-G**) Each dot represent a biological replicate. (**A-F**) Mean±SEM, * *p*<0.05, ** *p*<0.01, *** *p*<0.001, **** *p*<0.0001, ns *p*>0.1. *P* value is calculated by Student’s t-test, two-tailed, unpaired with Welch’s correction unless stated otherwise.

In a second *in vivo* model we performed the adoptive transfer of TCL1-355 into C57BL/6 mice as a model of chronic lymphocytic leukaemia (CLL) [48, 49]. In this model, leukemic B cells proliferate rapidly within the spleen which in theory should create areas depleted of glucose as within solid tumours which is what we are mimicking *in vitro* using Gal. Indeed, we have observed decreased glucose concentration in the spleens of CLL mice compared to healthy mice (Fig. 8F). Therefore, the CLL model can be employed as an *in vivo* model for glycolytic limitation. After the disease developed, mice were continuously infused via the tail vein with [U-^13^C] Glc tracer for 2 hours. When comparing B cells from healthy spleens against CLL B cells from diseased spleens, we observed that CLL cells significantly increased their relative flux from glucose towards serine and glycine while showing a reduced flux towards citrate, indicating low pyruvate oxidation (Fig. 8G). This altered metabolic phenotype between normal and leukemic B cells *in vivo* is partly recapitulated *in vitro* by switching cancer cells from Glc to Gal. In summary, by using Gal we are able to capture relevant metabolic features of tumour cells that occur in glucose depleted TME, specifically, the glycolytic rewiring towards serine to support cancer cell metabolism.

## Discussion

In the present study, we discover a novel rewiring of glycolytic flux towards SSP via a PKM2 block when the supply of glycolytic carbons is inadequate as modelled with Gal as a tool compound to mimic glycolytic limitation within the TME. In addition, we were able to corroborate previously described rewiring of glutamine metabolism towards SSP under glucose starvation. However, by identifying the block of lower glycolysis at PKM2, we were able to add another mechanistic layer that differentiates the glutamine flux under glycolytic limitation to either exclusively support SSP via PEPCK2 or TCA cycle via ME (Fig. 5H). We observed that both fluxes contribute to the survival and proliferation of several cancer cell lines and to supply the 1C cycle with the necessary carbons to perform biosynthetic, bioenergetic and redox balance functions. Cancer cells *in vitro* usually engage in aerobic glycolysis due to the ample availability of glucose. This does not reflect glucose levels and cancer cell metabolism within the TME. A key metabolite that drives the metabolic phenotype of cancer cells is glucose and its availability, which has been shown to be decreased in the tumour interstitial fluid (TIF) when compared to plasma [22, 23]. TIF measurements represent the average level of glucose across the area of the whole tumour. However, inside the tumour, areas do exist where cells are even more deprived of glucose (Fig. 8C) and those cells withstanding such severe metabolic stress (alongside other niche stresses) are the plastic cells that are capable of surviving the metastatic cascade [15]. Uncovering the metabolic rewiring in these deprived cells hold therapeutic promise to target cancer growth and metastasis [25].

By culturing cancer cells under glycolytic stress conditions, we were able to see their ability to use the same carbon source for different functions (metabolic plasticity) upon the PKM2 block and rewiring of carbons towards SSP. On the other hand, cancer cells also re-direct the glutamine flux towards SSP showcasing an example of metabolic flexibility [14]. PEPCK2 is the main instigator of glutamine flux rewiring towards SSP. It is intriguing that the mitochondrial isoform of PEPCK is involved under glycolytic limitations, as usually this isoform is only involved when cytoplasmic NADH levels are not depleted [50]. However, under glycolytic limitations, NAD/NADH ratio is increased and yet the cells still preferably engage PEPCK2 over cytoplasmic PEPCK1 whose involvement would transfer mitochondrial NADH to the cytoplasm via malate oxidation [50]. This finding can be explained by the possibility that the gluconeogenesis pathway is activated mainly to support SSP and therefore, cytoplasmic NADH is not required to reduce 1,3-bisphosphoglycerate into glyceraldehyde 3-phosphate as per the conventional gluconeogenesis pathway. Furthermore, in our system we did not observe a proliferative advantage in activating PEPCK2 under glycolytic limitation contrary to what has previously been published on PEPCK2. One major difference in our setup compared to earlier studies that could explain this observation is the usage of cell lines which have PHGDH amplification and are capable of increasing their serine synthesis flux independent of PEPCK2.

The diversion of the glycolytic flux away from lower glycolysis and towards SSP offers many metabolic advantages for the cell. For bioenergetics, SSP and subsequent 1C cycle generate two NADH and one ATP per 3-PG molecule, while lower glycolysis generates one ATP and consumes one NADH. Meaning if energy is the main constraint for the cell under glycolytic limitation, it will benefit more from running SSP and the subsequent serine catabolism and formate overflow [24]. Furthermore, the additional advantage of sustaining the flux towards serine catabolism lies in its pathway flexibility. The fate of the 1C can be tailored for the cell’s need; full oxidisation of 1C via ALDH1L2 will generate NADPH, the MTHFD1L reaction will generate ATP, or it can be incorporated in purines and dTMP in the cytosol [25]. Here we show that under glycolytic limitation, the rewiring towards SSP and 1C cycle supports all the three different pillars of metabolism, specifically the mitochondrial part of the 1C cycle (Fig. 7). In fact, we observe that even though the cells increase the expression of ALDH1L2 and subsequently lose more carbon as CO_2_, formate overflow is still increased, highlighting the non-biosynthetic benefit of running the mitochondrial branch of the 1C cycle under glycolytic limitation.

Cancer cells preferentially express PKM2 over other isozymes of pyruvate kinase as it provides them with the metabolic flexibility to regulate the glycolytic flux, as PKM2 can be regulated via allosteric signals that shifts its form between a high activity tetramer and a low activity dimer [42]. This allows cancer cells to rewire their glycolytic flux towards biosynthetic pathways when needed by expressing the dimer form of PKM2 to accommodate higher proliferative rates under high glycolytic flux [42]. In the present study, we observed a complete block PKM2 metabolic activity when the glycolytic flux is limited to rewire the limited carbon input towards SSP. Previous work has shown that serine acts as an allosteric activator of PKM2, while the lack of extracellular serine lead to an inhibition of PKM2 activity [51, 52]. Here we observed that when glucose is limited, even with serine/glycine available in the media, PKM2 glycolytic flux was inhibited. Indicating that even when serine was available for the cells, the lack of glycolytic carbons inhibited pyruvate kinase activity to rewire the glycolytic flux towards SSP. This observation highlights an intriguing difference between de novo serine synthesis flux and extracellular serine consumption and potentially could be explained by recent work that has shown that PHGDH functions beyond its catalytic activity within tumour cells and can allosterically regulate the function of phosphofructokinase [53]. Moreover, here we presented evidence that the removal of MTHFD2 or MTHFD1L was sufficient to partially re-establish PKM2 activity, meaning that mitochondrial 1C cycle is able to directly regulate PKM2 function alongside other regulators like F 1,6-BP. This would hint at a possible link between the PKM2 block and the increased formate overflow under glycolytic limitation. Entangling the direct cause of PKM2 block and the link with 1C cycle is the subject of ongoing work in our lab.

The findings presented here highlight the flexibility and plasticity of the biochemical network in tumour cells. These adaptive traits present a challenge in developing precision-medicine drugs that target the cells’ metabolism. For such therapeutic strategies to be successful, context dependent understanding of environmental conditions and nutrient availability of the TME need to be considered alongside specific oncogenic driver mutations [54].

## Acknowledgements

We are grateful to Stephanie Kreis (University of Luxembourg) for providing us with B16-F10, A375, SKMeL28 cells, Clement Thomas (LIH, Luxembourg) for providing 4T1 and BT-20 cells and Lewis Cantley (Harvard Medical School) for providing MDA-MB-468 WT and MTHFD1LKO cells. We are grateful for all the technical and analytical support from the different metabolomics platforms towards the work presented here and in specific: Francois Bernardin from the metabolomics platform at LIH, Xiangyi Dong, Floriane Gavotto and Lucia Gallucci from the LCSB metabolomics platform, Luxembourg. We are thankful for the support of the National Cytometry Platform (Quantitative Biology Unit, LIH) for support with flow cytometry measurements and analysis. Figures 1A, 1B, 3K, 5A, 5H, 7A and 8A were created with biorender.com.

M.B. is supported by the Fondation du Pélican de Mie et Pierre Hippert-Faber, under the aegis of the Fondation de Luxembourg. M.B. and J.M. are supported by the Luxembourg National Research Fund (FNR) ATTRACT program (A18/BM/11809970). J.M. is supported by FNR-PRIDE NEXTIMMUNE (PRIDE/11012546), FNR-PRIDE i2Tron (PRIDE19/14254520). E.L. is supported by the FNR-CORE program (C16/BM/11282028 and C20/BM/14591557), by a Proof of Concept FNR grant (PoC/18/12554295), a PRIDE17/11823097 and by i2Tron (PRIDE19/14254520). E.G., J.P., and E.M. are supported by grants from the FNR and Fondation Cancer (PRIDE15/10675146/CANBIO, C20/BM/14582635, and C20/BM/14592342). E.V. is supported by FNRS-Télévie (7.4509.20 and 7.4572.22).

## Author’s contribution

M.B. and J.M. conceptualised and designed the study. M.B. performed all of the experiments. A.O., E.V., E.G., M.S., C.P., J.P., E.M., S.P.N. and E.L., assisted for the design and performing of the *in vivo* work. L.N., N.I.L. and M.W.R. aided in the generation of different KO cell lines used. D.S., M.W., C.J. and A.L. performed YSI, LC-MS, GC-MS and IC-MS measurements while M.B. performed the corresponding data analysis according to D.S., M.W., C.J., A.L., J.M., instructions. Original draft written by M.B. and J.M. Review and editing done by all authors.

## Material and Methods

### Chemicals

Galactose, glucose, serine, glutamine and glycine were purchased from Sigma Aldrich and stock solutions were prepared in ^MQ^water. BI-4916 and TEPP-46 were purchased from MedChemExpress and stocks were prepared in DMSO. BI-4916 was always used at a final concentration of 15μM while TEPP-46 was used at 100μM.

### Cell culture

MDA-MB468, 4T1, MDA-MB-231, HCT116, LN-229 and B16-F10 were cultured in DMEM (Thermo Fisher Scientific, 1443001) supplemented with 17.5 mM glucose, 2mM glutamine and 10% FBS. BT20 were cultured in advanced DMEM/F12 (Thermo Fisher Scientific, 12634010) supplemented with 2 mM glutamine and 10% FBS. A375 and SKMeL28 were cultured in RPMI 1640 (Thermo Fisher Scientific, 61870010) supplemented with 10% FBS. All cells were maintained at 37°C and 5% CO_2_ and routinely tested for mycoplasma contamination. Serine/glycine starvation experiments were carried out using customised DMEM (Thermo Fisher Scientific, lot No. GME11933004) without phenol red, glucose, serine, glycine, tryptophan, pyruvate and glutamine with the missing ingredients added according to the concentrations present in full DMEM and experimental setup. All experiments (except anchorage independent growth assays, see below) were seeded on D-1 and cells were allowed to adhere and attach overnight. Medium was aspirated and experimental medium plus treatment was added for the indicated time points in figures.

MDA-MB-468 WT and MTHFD1L-KO were received from Lewis Cantley lab, Weill Cornell Medical College, USA [55]. STR profiling in our lab in 2021 authenticated MDA-MB-468. LN-229 WT and AMPK-dKO cells were obtained from Michael Ronellenfitsch, Dr Senckenberg Institute of Neurooncology, University Hospital Frankfurt, Goethe University and have been previously described in [46], B16-F10, A375 and SKMeL28 cells were received from Stephanie Kreis lab, University of Luxembourg, Luxembourg. 4T1 and BT20 were received from Clement Thomas lab, Luxembourg institute of Health, Luxembourg. MDA-MB-231 were obtained from Alexei Vasquez lab, Beatson Institute of Cancer Research, UK. All cell lines were originally purchased from ATCC.

### Lentiviral mediated knockdown of *PCK2*

The procedure followed was described in detail in [56]. For silencing *PCK2* gene in MDA-MB-468 cells, pGIPZ shPCK2-2: ATTATTGGACAGTCTTTGT (V3LHS_410309) vector plasmid purchased from Horizon Discovery was used.

### CRISPR/Cas9 KO of Pck2, MTHFD2, ALDH1L2

MDA-MB-468 cells harbouring deletion of MTHFD2 have been previously generated in [56]. To KO *Pck2* in 4T1 and B16-F10, we used the same procedure described in [56]; two different gRNAs were used: 1. TATGCGTATTATGACCCGCC (Vector ID: VB211125-1122kdb, gRNA #654) to generate 4T1 clone A9 and B16-F10 clone A5. 2. GCGGGTCATAATACGCATAC (Vector ID: VB211125-1123gfv, gRNA #653) to generate 4T1 clone B7 and B16-F10 clone A9. For *ALDH1L2* deletion in MDA-MB-468 cells the same procedure in [56] was utilised with two different gRNAs: 1. GCAGAAGCCTACAGATCCGT (Vector ID: VB190719-1044jbf, gRNA #1529) to generate MDA-MB-468 clone F11. 2. GCGCTCCGGCGCTTCTCCAC (Vector ID: VB190719-1054khg, gRNA #1^st^ Exon) to generate MDA-MB-468 clone D1.

### Animal experiments

All experiments involving laboratory animals were conducted in a specific pathogen-free animal facility with the approval of the animal welfare structure of LIH and the Luxembourg Ministry for Agriculture, Viticulture and rural development under the references LUPA 2021/08 and LUPA 2021/22. Mice were treated in accordance with the European Directive 2010/63/EU. C57BL/6 mice were purchased from Janviers Labs (France).

### TCL1-355 adoptive transfer

The TCL1-355 murine cell line was kindly provided by Dr Efremov [49]. A total of 2,000 TCL1-355 cells resuspended in DMEM without phenol red were injected intravenously (iv; 100 μL) in the tail vein of C57BL/6 mice. CLL progression was monitored by determining the percentage of CD5^+^CD19^+^ circulating CLL cells in the peripheral blood using flow cytometry. 14 days after iv injection, leukemic mice and age matched control mice were anaesthetised with 2.0 – 2.5% of isoflurane and continuously infused with [U-^13^C] glucose (50% solution in 0.9% NaCl) for 2h at a constant rate of 1.8g/Kg/h via the tail vein. At the end of the infusion, mice were euthanized via cervical dislocation and blood, organs and cells collection was rapidly performed.

Blood sampling was performed before and after infusion. Plasma was collected after two successive centrifugations at 1000 RCF for 10 min. B cells were isolated from the spleen. Briefly, suspension of splenocytes was prepared using gentleMACS™ Dissociator (Miltenyi). Then, B cells isolation (negative sorting) was performed using MojoSort™ Mouse Pan B Cell Isolation Kit II (Miltenyi), according to manufacturer’s instructions. Cells were counted prior to pelleting and snap freezing. Pieces of spleen were also snap frozen in liquid nitrogen for metabolomics.

### B16-F10 subcutaneous model

A total of 250,000 B16-F10 cells resuspended in PBS were injected subcutaneously (100 μL) into the flank of C57BL/6 mice and tumours were allowed to grow for 16 days for one group and 9 days for the second group. At endpoint, both groups were infused with [U-^13^C] glucose as described above. After infusion, mice were euthanized with cervical dislocation and blood was collected into heparin tubes to separate plasma. Plasma and tumour tissue were snap frozen in liquid nitrogen.

### Western blot

KOs and KDs cell pellets were directly obtained from cell culture by pelleting cells at 350 RCF for 5 min followed by a PBS wash and then freezing at −20°C awaiting cell lysis. Probing protein expression after treatments: cells were cultured at 150,000 cells/well density in a 6-well plate on D-1. At time point of extraction, the plate was placed on ice, medium was aspirated and cells were washed in ice-cold PBS. Cell lysis buffer (150 mM NaCl, 1 mM EDTA, 50 mM Tris-HCl, 1% NP-40 with proteases and phosphatases inhibitors (Roche)) was added directly to the wells and cells were scraped and collected in pre-cooled Eppendorf tubes followed by sonification for 10 min. Lysates were spun at 16,000 RCF, 4°C for 10 minutes, the supernatant was collected in new tubes and the concentration of protein was determined with a Bradford assay. 20 – 40 μg of total protein was loaded into 15-well or 12-well RunBlue 4-12% Bis-Tris gels (Westburg) with the addition of 10mM DTT (Sigma Aldrich) and 4x NuPage LDS Sample buffer (Thermo Fisher Scientific) and gels were run for 90 minutes at 120V. Protein bands were blotted onto nitrocellulose membrane for 2h at 30 V. Membranes were blocked using 5% milk powder in TBS-0.1% Tween solution for 1h at RT. Membranes were incubated in the corresponding primary antibody overnight at 4°C and for 1h at RT in secondary antibodies. Detection of protein bands was done using Odyssey CLx Infrared Imaging system (LI-COR). Image StudioLite software Vers.5.2 (LI-COR) was used for quantifications and analysis. Primary antibodies used in this paper: AMPKα (Cell Signalling Technology, 5831S), p-AMPK^Thr172^ (Cell Signalling Technology, 72335S), ACC (Cell Signalling Technology, 3676S), p-ACC^Ser79^ (Cell Signalling Technology, 11818S) MTHFD2 (Cell Signalling Technology, ab151447), ALHD1L2 (Sigma Aldrich, HPA020549 and Proteintech, 21391-1-AP), β-Actin (Cell Signalling Technology, 3700), PSAT1 (Sigma Aldrich, HPA042924), PSPH (Sigma Aldrich, HPA020376), PHGDH (Sigma Aldrich, HPA021241), PEPCK2 (Cell Signalling Technology, 6924S) and MTHFD1L (Proteintech, 16113-1-AP). All antibodies were used at 1:1000 dilution unless stated otherwise. Secondary antibodies: IRDye 680RD Goat Anti-Mouse IgG (H + L) (926-68070) and IRDye 800CW Donkey Anti-Rabbit IgG (H + L) (926-32213) from LI-COR (used in a 1:10,000 dilution)..

### Flow cytometry analysis of Apoptosis

250,000 cells (MDA-MB-468, B16-F10 and A375) or 200,000 cells (4T1) were seeded in 6-well plate, allowed to attach overnight and treated as indicated for 48h. At endpoint, medium was collected, cells were washed with PBS and PBS was collected followed by cell trypsinisation. Cells were pelleted and washed with ice-cold PBS. Each sample was stained for 15 min on ice and in the dark with 50 μL AnnexinV staining solution (5% AnnexinV-FITC antibody in AnnexinV binding buffer (10 mM HEPES pH 7.4, 140 mM NaCl, 2.5 mM CaCl_2_, 0.1% BSA in ^MQ^water)). Directly prior to measurements, 450 μL of propidium iodide (PI) staining solution was added (11.1 μg/ml in AnnexinV binding buffer). Measurements were done using NovoCyte Quanteon (Agilent) and software NovoExpress (version 1.5.0). FlowJo version 10.6.2 was used for analysis.

### Proliferation assays

Cells were seeded at a concentration of 25,000 cells/well in a 12-well plate or 12,500 cells/well in a 24-well plate in triplicates. Separate triplicate wells were counted when treatment was added to set 0h count for fold change calculations. Medium and treatment were replaced at 48h and 72h. At endpoint, cells were trypsinised and viable cell count was obtained using Countess^™^ automated counter (Thermo Fisher). Growth rate in Fig. 6A was calculated using the equation: [*ln*(final cell count/initial cell count)]/duration.

### Anchorage Independent 3D growth assays

Cells were cultured in ultra-low attachment flask (Greiner) for a minimum of five days. Cells were collected and spun at 100 RCF for 5 min and afterwards resuspended in Ca^+2^ and Mg^+2^ free PBS. 5,000 cells were seeded in 96-well ultra-low attachment plates (S-Bio or facellitate) already in the experimental medium and the plates were spun at 100 RCF for 10 minutes. 3D growth was determined using IncuCyte® S3 Live-Cell Analysis system (Sartorius) over the indicated time periods.

### RNA extraction, cDNA synthesis and RT-qPCR

300,000 cells were seeded in 6-well plate and treated for 24h as indictaed in the figures. RNA was extracted directly from the plates using RNeasy Mini Kit (Qiagen, 74104). 2μg of RNA were used for cDNA synthesis using High Capacity cDNA Reverse Transcription Kit (Thermo Fischer Scientific, 4368814). For qPCR, we used Fast SYBR™ Green Master Mix (Thermo Fischer Scientific, 4368814) with 20 ng cDNA/sample in triplicates or duplicates and the reaction was conducted at 95°C for 20 s followed with 40 cycles of 95°C for 1 s and 60°C for 20 s. The QuantStudio 5 Real-Time PCR System was used (Applied Biosciences, ThermoFisher Scientific). CT values were determined by QuantStudio Design and Analysis v1.5.1 software (Applied Biosciences, ThermoFisher Scientific). Fold change of mRNA expression was determined using ddCt algorithm. Primers used in this study are listed in Table S1.

**Supplementary Table 1.**
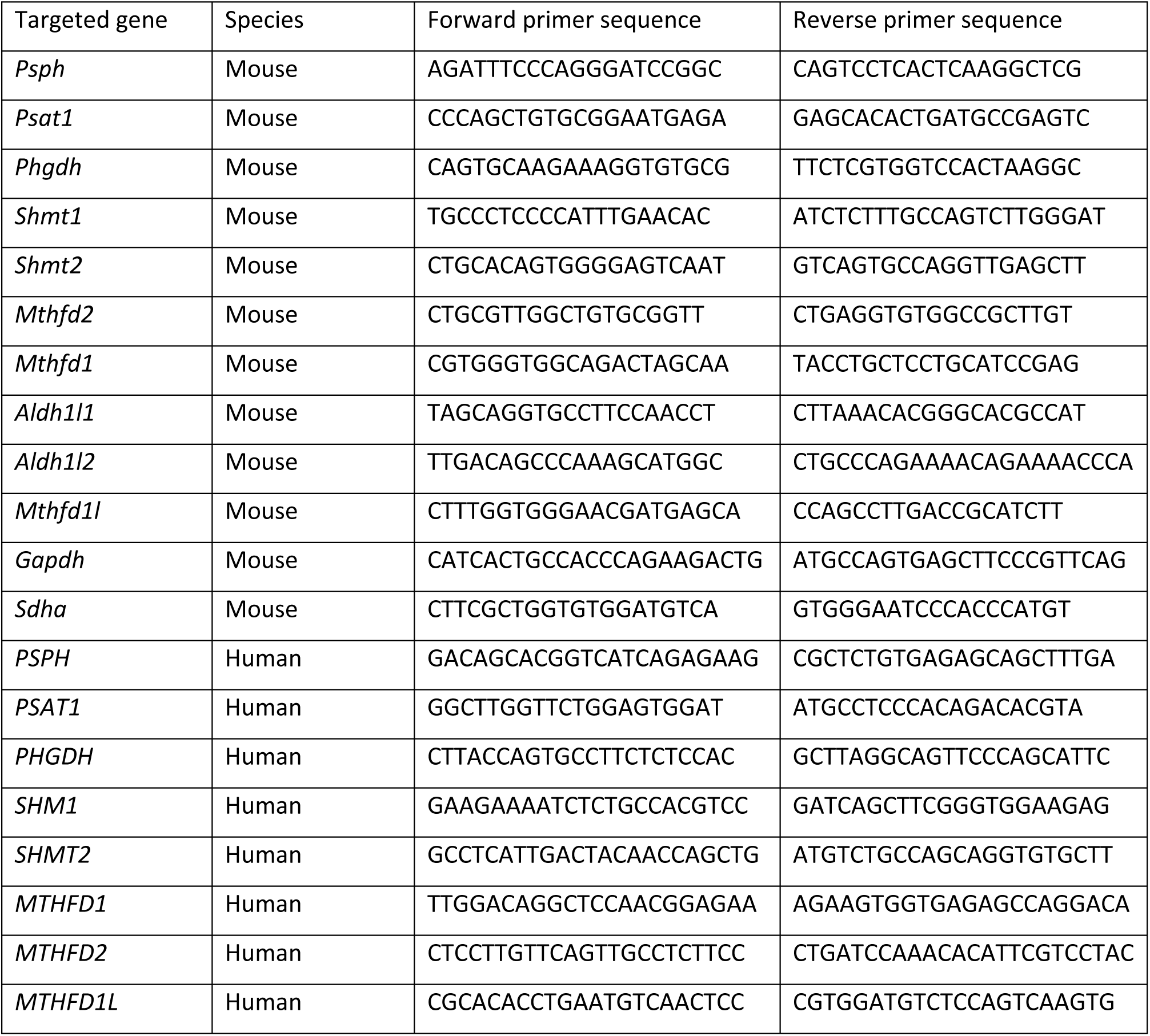
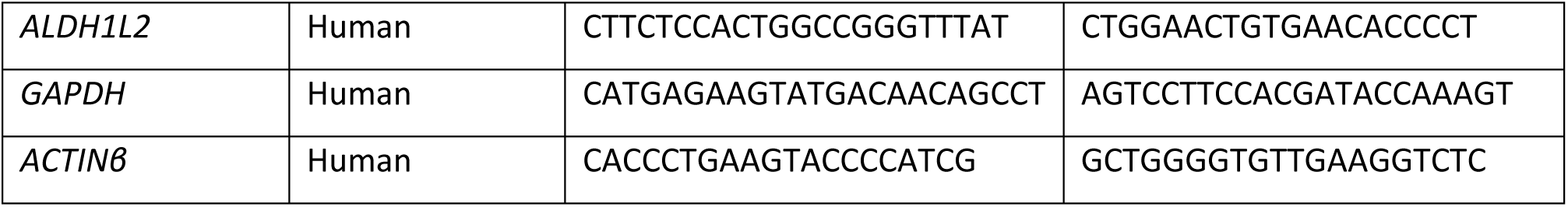
List of primers used in this study and their sequences.

### Metabolomics

#### Stable Isotope Tracing and Metabolite Extraction

##### *In vitro* tracing and metabolite extraction

Stable isotope tracing experiments with [U-^13^C] glucose, [U-^13^C] galactose, [U-^13^C] glutamine, [U-^13^C] serine (Cambridge Isotope Laboratories, CLM-1396, CLM-1570, CLM-1822 and CLM-1574 respectively) and [U-^13^C] lactate (Sigma Aldrich, 485926) were performed in DMEM (Thermo Fisher Scientific, 1443001), RPMI 1640 for SILAC (Thermo Fisher Scientific, 88365), Advanced DMEM/F12 for SILAC (Thermo Fisher Scientific, A2494301) or customised DMEM (Thermo Fisher Scientific, lot No. GME11933004) depending on the cell lines used. These tracer mediums were supplemented with the missing metabolites according to the experimental setup to mimic the original formula of DMEM, RPMI 1640 or advanced DMEM/F12. For sodium [U-^13^C] lactate 5 mM final concentration was used. For glucose and galactose tracing 10% normal FBS was used while for the rest we used 10% dialysed FBS. Experimental set-up and extraction of metabolites for GC-MS analysis was done as described in [56]. Metabolite extraction for LC-MS was done as described in [24]. Metabolite extraction for IC-MS was identical to LC-MS.

##### *In vivo* metabolite extraction

Metabolite extraction from tumours, spleen and plasma was performed as detailed in [24]. The absolute quantification of glucose in plasma and spleen was performed using standard addition method. In brief, known concentrations of glucose were added serially to metabolite extracts of each sample. After measurements, a calibration curve was used to determine the concentration of glucose in the original samples by determining the x-axis intercept.

For B cells separated from the spleen, cells were washed with ice-cold PBS and subsequently 40 μL/one million cells extraction buffer [acetonitrile/^MQ^H_2_O/methanol (ratio, 3:2:5); liquid chromatography–MS (LC-MS) grade solvents)] was added to each cell pellet. Cells were mixed for 10 min on a thermomixer at 4°C at maximum speed, and then the tubes were centrifuged for 10 min at 16,100 RCF at 4°C. 50 µL of the supernatant was collected and transferred to an already-cooled LC-MS glass vial with inserts and stored at - 80°C until measurement.

#### YSI measurements and medium exchange rates

Medium samples were filtered (PVDF, 0.22 µm) prior measurement to remove particulates.

Absolute quantitative values for lactic acid and glutamine were acquired using a YSI 2950D Biochemistry Analyzer (Kreienbaum KWM). The instrument was calibrated and prepared according to the manufacturers’ instruction. For a precise and reliable quantification, external concentration curves of each target compound were prepared and measured in triplicates. Absolute uptake and release rates were calculated as described in [24].

#### GC-MS measurements

##### Formate measurements

Formate was derivatized to benzyl formate from the collected medium samples using the MCF protocol described in [24]. GC-MS analysis was performed using an Agilent 7890 A GC coupled to an Agilent 5975 C inert XL Mass Selective Detector (Agilent Technologies). A sample volume of 1 μL was injected into a Split/Splitless inlet, operating in split mode (20:1) at 280 °C. The gas chromatograph was equipped with a 30 m (I.D. 250 μm, film 0.25 μm) ZB-5MSplus capillary column (Phenomenex, 7HG-G030-11-GGA) plus 5 m GUARDIAN in front of the analytical column. Helium was used as carrier gas with a constant flow rate of 1.4 ml/min. GC oven temperature was held at 90 °C for 1 min and increased to 115 °C at 10 °C/min followed by 20 °C/min to 160 °C and a post run time of 4.25 min at 325 °C. Total run time was 10 min. Transfer line temperature was set to 280 °C. Mass selective detector (MSD) was operating under electron ionization at 70 eV. MS source was held at 230 °C and the quadrupole at 150 °C. For precise quantification, measurements were performed in selected ion monitoring mode. Target ions (m/z) and dwell times are listed below in Table S2. GC-MS chromatograms were processed using Agilent MassHunter Quantitative Analysis for GC-MS, Version B.08.00. Natural isotope correction and analysis of formate release rates was done as previously described in [24].

**Supplementary Table 2.** Dwell times and quantification ions for the derivatized formate.

| Derivative (Metabolite) | Quant-Ion (m/z) | Dwell Time (ms) |
| --- | --- | --- |
| Benzyl Formate M<sub>0</sub> | 136.1 | 40 |
| Benzyl Formate M<sub>1</sub> | 137.1 | 40 |
| IS-Benzyl Formate (<sup>13</sup>C,D) M<sub>0</sub> | 138.1 | 40 |
| IS-Benzyl Formate (<sup>13</sup>C,D) M<sub>1</sub> | 139.1 | 40 |
| IS-Benzyl Formate(<sup>13</sup>C,D) M<sub>2</sub> | 140.1 | 40 |

##### Intracellular metabolites measurements (Fig. S3A, C)

First, dried polar extracts were derivatized for 90 min at 55 °C with 20 μl of methoxyamine (c = 20 mg/ml) in pyridine under continuous shaking and subsequently for 60 min at 55 °C with additional 20 μl of MTBSTFA w/ 1% TBDMCS. GC-MS analysis was performed using an Agilent 7890B GC coupled to an Agilent 5977 A Mass Selective Detector (Agilent Technologies). A sample volume of 1 μl was injected into a Split/Splitless inlet, operating in splitless mode at 270 °C. Gas chromatograph was equipped with a 30 m (I.D. 250 μm, film 0.25 μm) ZB-35MS capillary column with 5 m guard column (Phenomenex). Helium was used as carrier gas with a constant flow rate of 1.2 ml/min. GC oven temperature was held at 100°C for 2 min and increased to 300°C at 10°C/min and held for 4 min. Total run time was 26 min. Transfer line temperature was set to 280°C. Mass selective detector (MSD) was operating under electron ionization at 70 eV. MS source was held at 230°C and the quadrupole at 150 °C. For precise quantification of the MID, measurements were performed in selected ion monitoring mode. The derivative measured here was pyruvic acid 1MeOX 1TBDMS with target ions 174.0 – 180.1 [M-57 fragment] and dwell times of 15 ms. The MetaboliteDetector software package (Version 3.220180913) was used for mass spectrometric data post processing, quantification, MID calculations, correction of natural isotope abundance, and determinations of fractional carbon contributions [57].

#### **IC-MS measurements** (Fig. 1C-I, Fig. S4D and S6I-K)

For the IC-MS measurements, polar extracts were analyzed as previously described [58] with the following adjustments. 50-75% of the total polar extract was dried overnight under vacuum. Dried polar metabolite pellets were reconstituted in 25 µl of water (Milli-Q Advantage A10) of which 20 µl was injected. For the ICMS system, we interfaced a Dionex ICS-6000+ ion chromatograph coupled with a QExactive high resolution mass spectrometer (Thermo Fisher Scientific). The guard, column, gradient settings, and suppressor type remained unchanged [58]. The electrospray ionization (ESI) was set as follows: spray voltage of 2.500 kV, auxiliary gas temperature of 420 °C, seep gas flow rate of 0, auxiliary gas flow rate of 15 units, and sheath gas flow rate of 50 units. The settings for acquisition of MS1 data were: m/z scan range of 80-750, resolution of 17500 at 200 m/z, automatic gain control (AGC) target of 1e6, and maximum injection time of 100 milliseconds. Data were acquired using two micro-scans.

#### LC-MS Measurements

Metabolite analyses were performed using a Vanquish Flex LC coupled to a Q Exactive HF mass spectrometer (Thermo Scientific). Chromatography was carried out with a SeQuant ZIC-pHILIC 5 µm particles column (150 × 2.1 mm) protected by SeQuant ZIC-pHILIC Guard (20 × 2.1 mm) pre-column. Column temperature was maintained at 45°C. The flow rate was set to 0.2 ml/min and the mobile phases consisted of 20 mmol/L ammonium carbonate in water, pH 9.2 (Eluent A) and Acetonitrile (Eluent B). The gradient was: 0 min, 80% B; 2 min, 80% B; 17 min, 20% B; 18 min 20% B; 19 min 80% B; 20 min 80% B (0.4ml/min); 24 min 80% B (0.4ml/min); 24.5 min 80% B. The injection volume was 5 µL. All MS experiments were performed using electrospray ionization with polarity switching enabled. The source parameters were applied as follows: sheath gas flow rate, 25; aux gas flow rate, 15; sweep gas flow rate, 0; spray voltage, 4.5 kV (+) / 3.5 kV (-); capillary temperature, 325°C; S-lense RF level, 50; aux gas heater temperature, 50°C. The Orbitrap mass analyzer was operated at a resolving power of 30,000 at 200 m/z in full-scan mode (scan range: *m/z* 75…1000; automatic gain control target: 1e6; maximum injection time: 250 ms).

Data from IC-MS and LC-MS measurements were acquired with Thermo Xcalibur software (Version 4.3.73.11) and analyzed with TraceFinder (Version 4.1). Natural isotope subtraction and subsequent analysis were performed as previously described using in-house scripts [24].

#### Oxygen consumption rate measurements

10,000 cells (MDA-MB-468, 4T1 and MDA-MB-231) or 20,000 cells (HCT116) were seeded into 96-seahorse microplate (Agilent) on D-1. After attachment, glucose and galactose medium were added and cells were incubated for 24h. Basal OCR was measured using XF96 Extracellular Flux Analyzer (Seahorse bioscience, Agilent) following the manufacturer’s protocol, Wave 2.6.0 software was used for analysis. OCR was normalised by the protein concentration of the corresponding wells following the protocol described in[59] and using a Bradford assay.

#### Statistics

Unless stated otherwise, unpaired two-tailed student’s t-test with Welch’s correction was used throughout the presented study as statistical tests were always performed between two groups. GraphPad PRISM (9.5.0) was used to calculate the *p* value. In the present study, *in vitro n* is defined as one independent experiment with at least two replicate wells (except for protein lysates and RNA samples where single wells were used). For *in vivo* experiments, *n* is defined as a sample from one mouse.

---

**Supplementary Figure 1.**
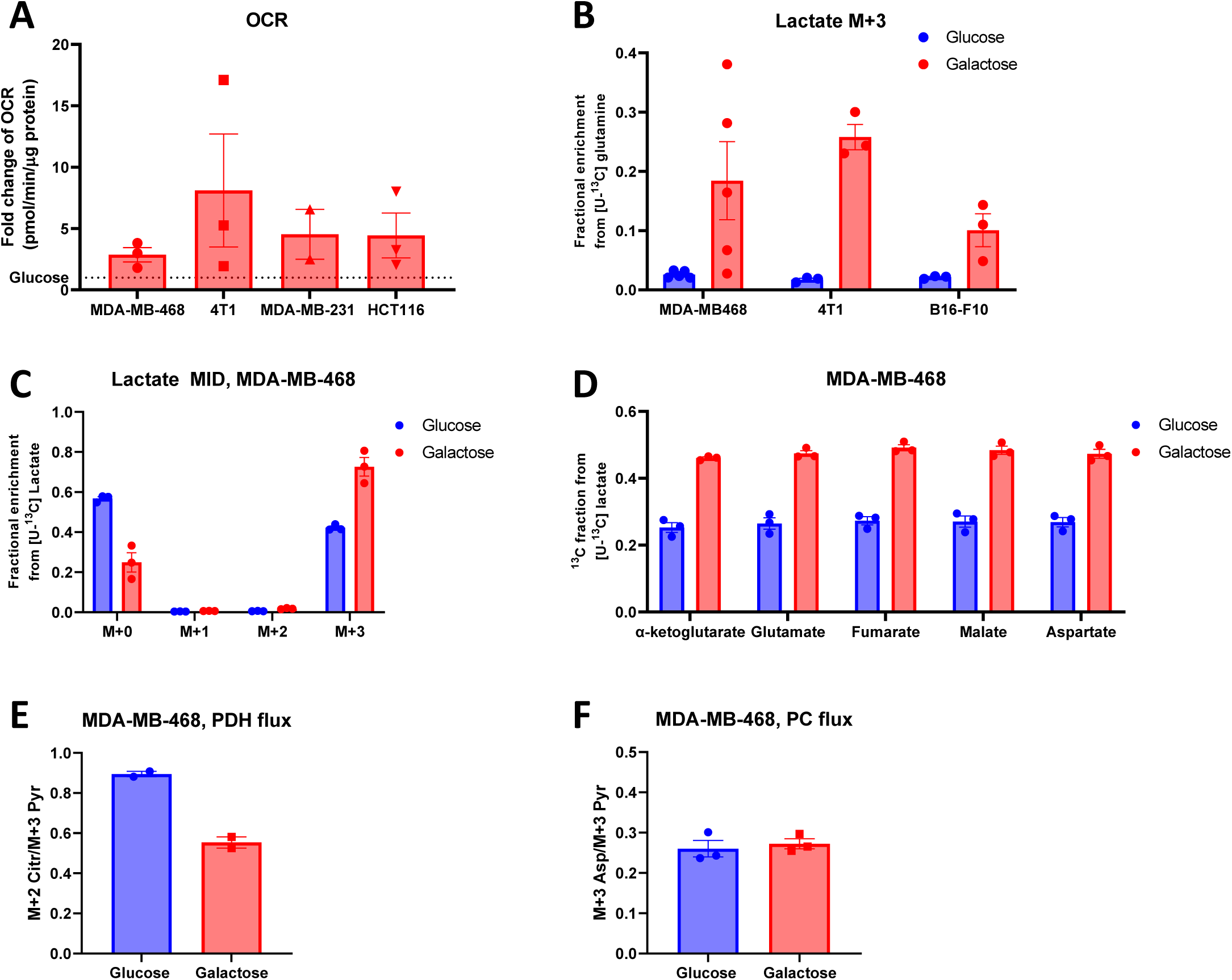
**A** Basal oxygen consumption rate (OCR) measured by seahorse in 4 different cancer cell lines after 24h incubation with galactose and presented as fold change to the same measurements with glucose. **B** Fractional enrichment in M+3 lactate from [U-^13^C] glutamine after 24h incubation with either glucose or galactose medium in MDA-MB-468, 4T1 and B16-F10. **C** Fractional enrichment in lactate from [U-^13^C] sodium lactate after 24h incubation with either glucose or galactose medium in MDA-MB-468. **D** ^13^C fraction in TCA cycle metabolites from [U-^13^C] sodium lactate after 24h incubation with either glucose or galactose medium in MDA-MB-468. **E** Pyruvate dehydrogenase (PDH) enzyme flux ratio calculated as M+2 citrate/M+3 pyruvate isotopologues from [U-^13^C] sodium lactate after 24h incubation with either glucose or galactose medium in MDA-MB-468. **F** Pyruvate carboxylase (PC) enzyme flux ratio calculated as M+3 aspartate/M+3 pyruvate isotopologues from [U-^13^C] sodium lactate after 24h incubation with either glucose or galactose medium in MDA-MB-468. (**A-F**) Each dot represents an independent experiment averaged from 3 different replicate wells. (**A-F**) Mean±SEM, * *p*<0.05, ** *p*<0.01, *** *p*<0.001, **** *p*<0.0001, ns *p*>0.1. *P* value is calculated by Student’s t-test, two-tailed, unpaired with Welch’s correction.

**Supplementary Figure 2.**
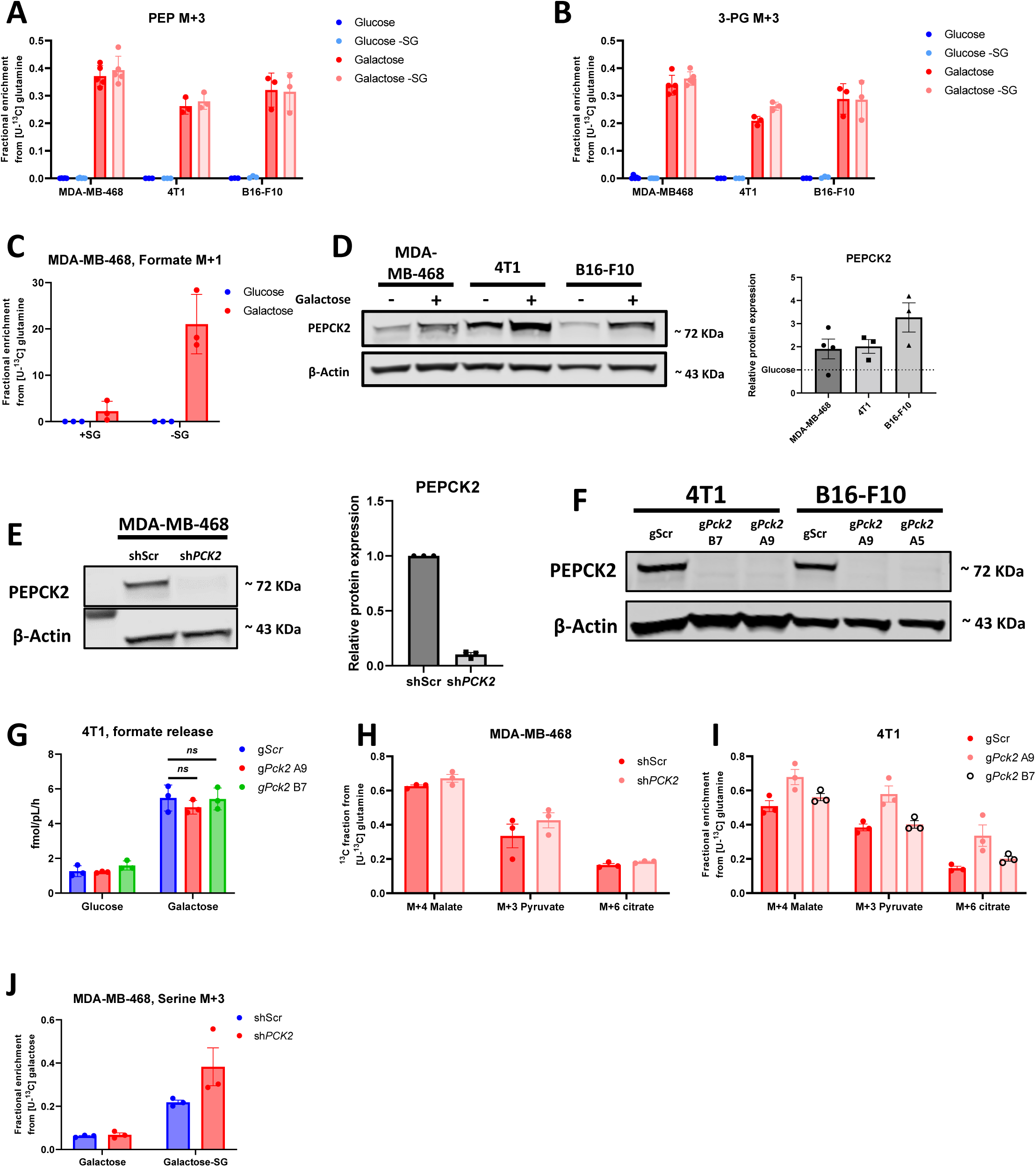
**A-B** Fractional enrichment in M+3 PEP **(A)** and M+3 3-PG **(B)** from [U-^13^C] glutamine after 24h incubation with either glucose ± serine and glycine (SG) or galactose ± SG medium in MDA-MB-468, 4T1 and B16-F10. **C** Fractional enrichment in M+1 formate from [U-^13^C] glutamine after 24h incubation with either glucose ± serine and glycine (SG) or galactose ± SG medium in MDA-MB-468. **D** Protein expression of PEPCK2 after 48h incubation with either glucose or galactose, with the corresponding quantification of signal intensities relative to β-actin as a loading control and presented as a fold change of expression under galactose to that of glucose. **E-F** Protein expression of PEPCK2 comparing scramble to PEPCK2 KD/KOs, with the corresponding quantification of these signals relative to β-actin as a loading control. **G** Absolute quantification of formate release from 4T1 after 24h incubation with either glucose or galactose. **H-I** Fractional enrichment in M+4 malate, M+3 pyruvate and M+6 citrate from [U-^13^C] glutamine after 24h incubation with galactose + SG medium and comparison between control and MDA-MB-468 PEPCK2 KD **(H)** and 4T1 PEPCK2 KOs **(I)**. **J** Fractional enrichment in M+3 serine from [U-^13^C] galactose after 24h incubation with galactose ± SG medium in MDA-MB-468 shScramble and sh*PCK2*. (**A-J**) Each dot represents an independent experiment averaged from 3 different replicate wells replicate unless stated otherwise. (**A-J**) Mean±SEM, * *p*<0.05, ** *p*<0.01, *** *p*<0.001, **** *p*<0.0001, ns *p*>0.1. *P* value is calculated by Student’s t-test, two-tailed, unpaired with Welch’s correction unless stated otherwise.

**Supplementary Figure 3.**
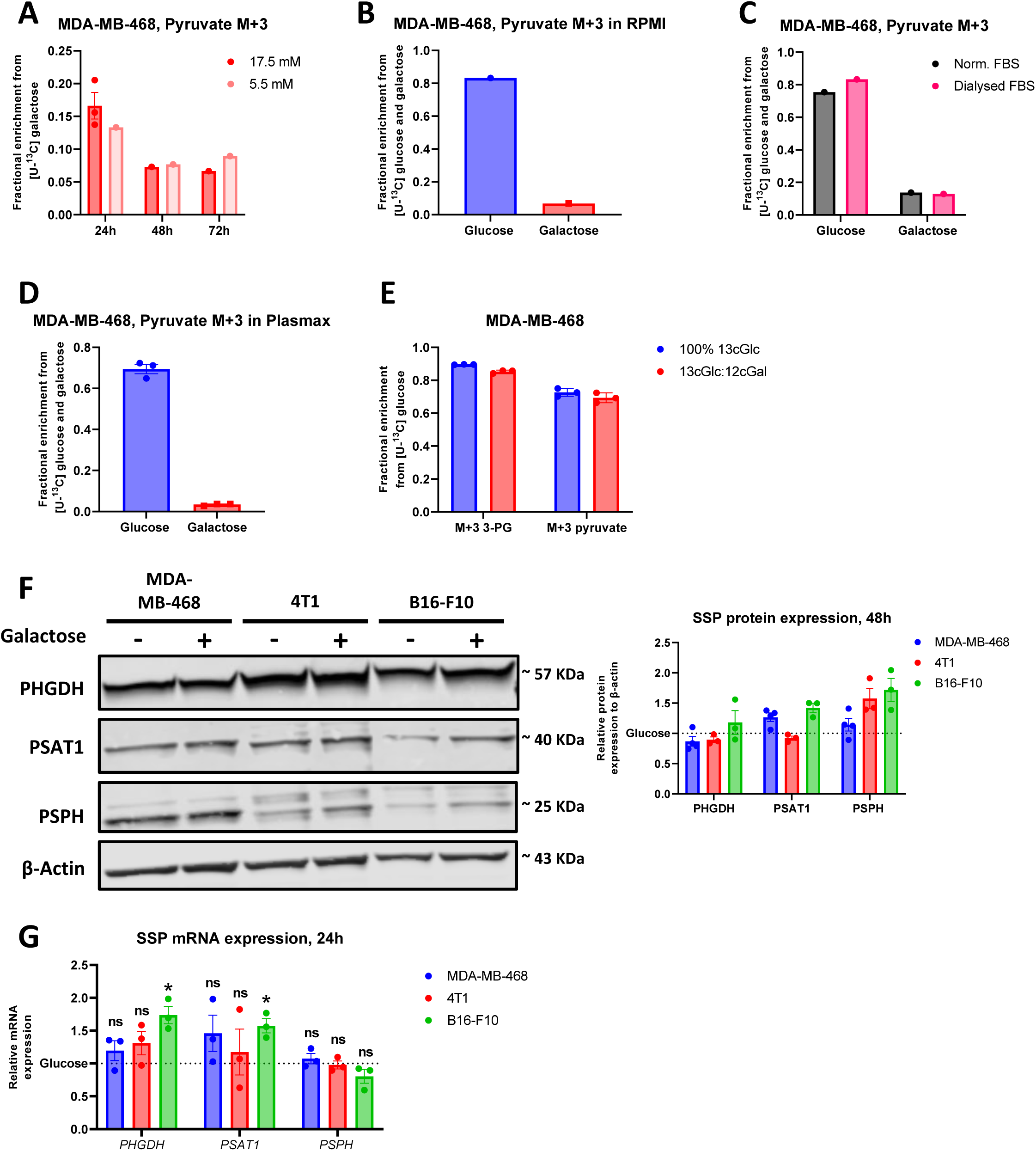
**A** Fractional enrichment into M+3 pyruvate using [U-^13^C] galactose after 24, 48, and 72 hour incubation and using 17.5 or 5.5 mM of the tracer in MDA-MB-468. **B** Fractional enrichment into M+3 pyruvate using [U-^13^C] glucose and galactose in RPMI1640 in MDA-MB-468 with 10% dialysed FBS. **C** Fractional enrichment into M+3 pyruvate using [U-^13^C] glucose and galactose after 24 hour incubation and using 17.5 mM of the tracer in MDA-MB-468 with 10% FBS or 10% dialysed FBS. **D** Fractional enrichment into M+3 pyruvate using [U-^13^C] glucose and galactose in physiological medium Plasmax in MDA-MB-468. **E** Fractional enrichment from 17.5 mM [U-^13^C] glucose into M+3 3PG and M+3 pyruvate after 24h incubation measured with LC-MS. **F** Protein expression of PHGDH, PSAT and PSPH after 48h incubation with either glucose or galactose, with the quantification of signal intensities relative to β-actin as a loading control and shown as a fold change of expression under galactose to that of glucose. **G** mRNA expression of *PHGDH*, *PSAT1*, *PSPH* relative to *GAPDH* and *ACTINβ* or *Gapdh and Sdha* after 24h incubation with either glucose or galactose measured with RT-qPCR. Data presented as fold change to the mRNA expression in cells cultured with glucose. (**A**-**G**) Each dot represent an independent experiment (**A-E** each independent experiment is from at least 3 different replicate wells). (**A**-**G**) Mean±SEM, * *p*<0.05, ** *p*<0.01, *** *p*<0.001, **** *p*<0.0001, ns *p*>0.1. *P* value is calculated by Student’s t-test, two-tailed, unpaired with Welch’s correction.

**Supplementary Figure 4.**
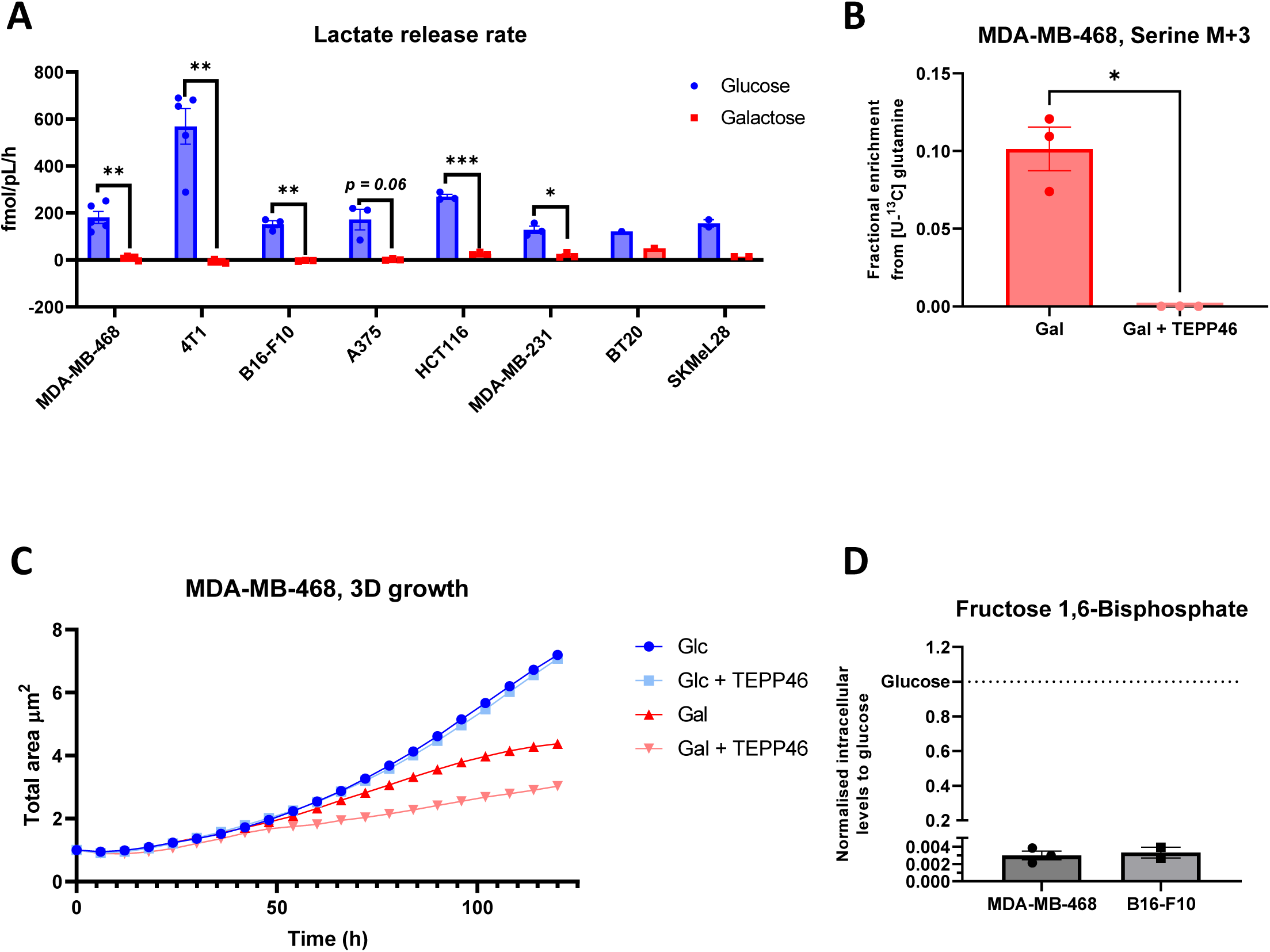
**A** Absolute quantification of lactate released into the medium per hour of incubation with normalisation to packed cell volume per condition in either glucose or galactose in multiple cancer cell lines. **B** Fractional enrichment from [U-^13^C] glutamine into M+3 serine after 24h incubation measured with LC-MS. **C** 3D growth of MDA-MB-468 aggregates using anchorage-independent conditions when treated with 100 µM of TEPP-46 with either glucose or galactose. 1 independent experiment is represented with at least 6 different replicate wells. **D** Intracellular levels of fructose 1,6-BP under galactose represented as a fold change of the levels under glucose in MDA-MB-468 and B16-F10. (**A**, **B** and **D**) Each dot represent an independent experiment (from 3 different replicate wells). (**A**-**C**) Mean±SEM, * *p*<0.05, ** *p*<0.01, *** *p*<0.001, **** *p*<0.0001, ns *p*>0.1. *P* value is calculated by Student’s t-test, two-tailed, unpaired with Welch’s correction.

**Supplementary Figure 5.**
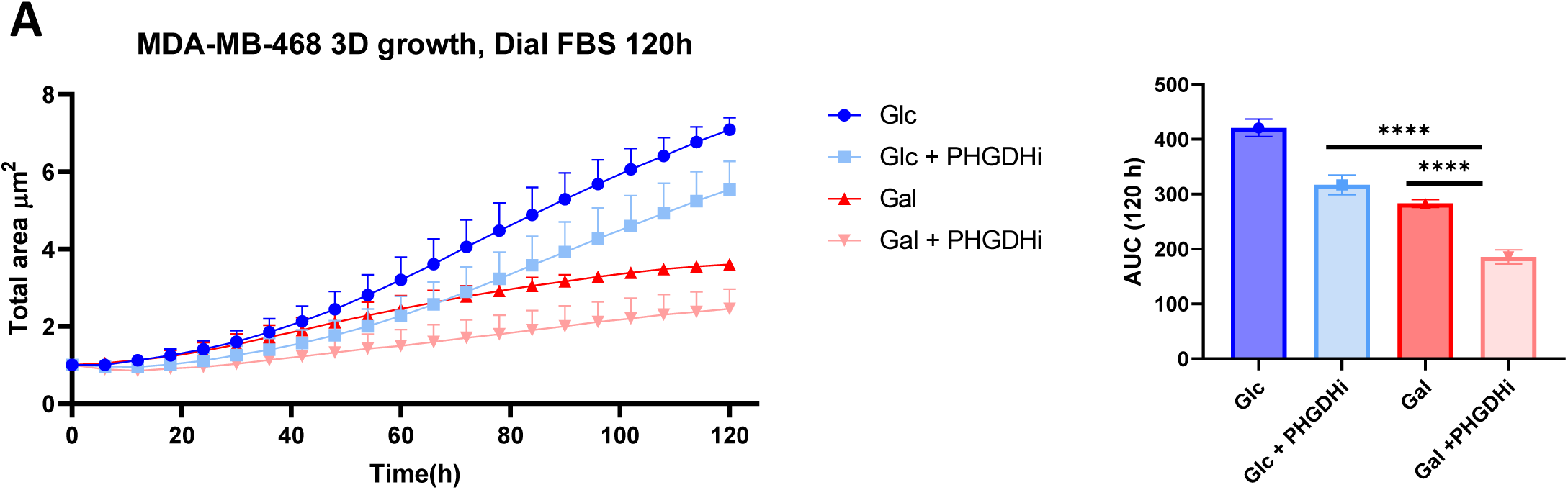
**A** 3D growth of MDA-MB-468 aggregates using anchorage-independent conditions when treated with 15 µM of PHGDHi with either glucose or galactose with 10% dialysed FBS, with the corresponding AUC. 3 independent experiments are represented each with at least 6 replicate wells. Mean±SEM, * *p*<0.05, ** *p*<0.01, *** *p*<0.001, **** *p*<0.0001, ns *p*>0.1. *P* value is calculated by Student’s t-test, two-tailed, unpaired with Welch’s correction.

**Supplementary Figure 6.**
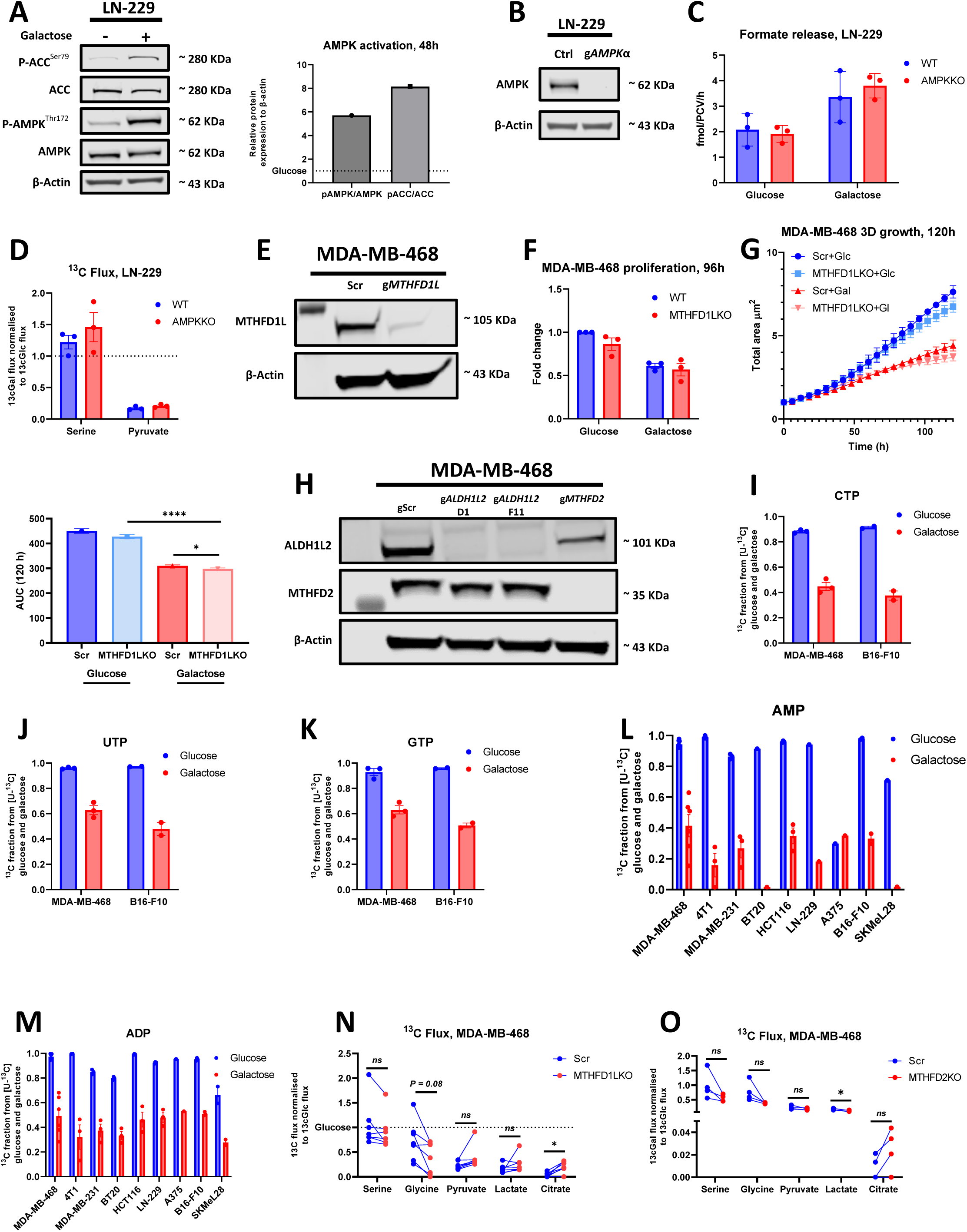
**A** Protein expression of p-AMPK, AMPK, p-ACC and ACC in LN-229 after 48h incubation with either glucose or galactose, with the quantification of signal intensities relative to β-actin as a loading control and shown as a fold change of expression under galactose to that of glucose. **B** Protein expression of AMPK comparing control to the corresponding KO with β-actin as a loading control. **C** Absolute quantification of formate release from LN-229 control and AMPKKO after 24h incubation with either glucose or galactose. **D** The normalised relative ^13^C flux under galactose towards serine and pyruvate is represented as a fold change of the same ratio under glucose in LN-229 control and AMPKKO. **E** Protein expression of MTHFD1L comparing scramble to the corresponding KO with β-actin as a loading control. **F** 2D proliferation of MDA-MB-468 Scramble and MTHFD1LKO cells cultured in either glucose or galactose as measured with cell counts at 96h relative to proliferation of scramble cells under glucose. **G** 3D growth of MDA-MB-468 control and MTHFD1LKO cell aggregates using anchorage-independent conditions in either glucose or galactose, with the corresponding AUC. 3 independent experiments are represented each with at least 6 different replicate wells. **H** Protein expression of MTHFD2 and ALDH1L2 comparing scramble to the corresponding KOs with β-actin as a loading control. **I-K** Fractional enrichment in CTP **(I)**, UTP **(J)** and GTP **(K)** from [U-^13^C] glucose and galactose after 24h incubation. Data shown is the mean±SEM of 3 (MDA-MB-468) or 2 (B16-F10) independent experiments each being the average of 3 different replicate wells. **L-M** ^13^C fraction in AMP **(L)** and ADP **(M)** from [U-^13^C] glucose and galactose after 24h incubation in multiple cell lines. **N**-**O** The normalised relative ^13^C flux under galactose towards multiple metabolites represented as a fold change of the same ratio under glucose in MDA-MB-468 scramble compared to MTHFD1LKO (**N**) and MTHFD2KO (**O**). (**A, C-D, F and I-O**) each dot represent an independent experiment averaged from 3 different replicate wells unless stated otherwise. (**A-O**) Mean±SEM, * *p*<0.05, ** *p*<0.01, *** *p*<0.001, **** *p*<0.0001, ns *p*>0.1. *P* value is calculated by Student’s t-test, two-tailed, unpaired with Welch’s correction unless stated otherwise.

## References

1. Crabtree, H.G., Observations on the carbohydrate metabolism of tumours. Biochem J, 1929. 23(3): p. 536–45.

2. Hanahan, D. and R.A. Weinberg, Hallmarks of cancer: the next generation. Cell, 2011. 144(5): p. 646–74.

3. Warburg, O., The Metabolism of Carcinoma Cells1. The Journal of Cancer Research, 1925. 9(1): p. 148–163.

4. Vander Heiden, M.G., L.C. Cantley, and C.B. Thompson, Understanding the Warburg effect: the metabolic requirements of cell proliferation. Science, 2009. 324(5930): p. 1029–33.

5. Amendola, C.R., et al., KRAS4A directly regulates hexokinase 1. Nature, 2019. 576(7787): p. 482–486.

6. Fernandez-de-Cossio-Diaz, J. and A. Vazquez, Limits of aerobic metabolism in cancer cells. Sci Rep, 2017. 7(1): p. 13488.

7. Locasale, J.W., et al., Phosphoglycerate dehydrogenase diverts glycolytic flux and contributes to oncogenesis. Nat Genet, 2011. 43(9): p. 869–74.

8. Osthus, R.C., et al., Deregulation of glucose transporter 1 and glycolytic gene expression by c-Myc. J Biol Chem, 2000. 275(29): p. 21797–800.

9. Vazquez, A., et al., Catabolic efficiency of aerobic glycolysis: The Warburg effect revisited. BMC Systems Biology, 2010. 4(1): p. 58.

10. Ying, H., et al., Oncogenic Kras maintains pancreatic tumors through regulation of anabolic glucose metabolism. Cell, 2012. 149(3): p. 656–70.

11. de Alteriis, E., et al., Revisiting the Crabtree/Warburg effect in a dynamic perspective: a fitness advantage against sugar-induced cell death. Cell Cycle, 2018. 17(6): p. 688–701.

12. Luengo, A., et al., Increased demand for NAD(+) relative to ATP drives aerobic glycolysis. Mol Cell, 2021. 81(4): p. 691–707.e6.

13. Wang, Y., et al., Saturation of the mitochondrial NADH shuttles drives aerobic glycolysis in proliferating cells. Molecular Cell, 2022. 82(17): p. 3270–3283.e9.

14. Bergers, G. and S.M. Fendt, The metabolism of cancer cells during metastasis. Nat Rev Cancer, 2021. 21(3): p. 162–180.

15. Fendt, S.-M., C. Frezza, and A. Erez, Targeting Metabolic Plasticity and Flexibility Dynamics for Cancer Therapy. Cancer Discovery, 2020. 10(12): p. 1797–1807.

16. Cantor, J.R., The Rise of Physiologic Media. Trends in Cell Biology, 2019. 29(11): p. 854–861.

17. Díaz-Ruiz, R., et al., Mitochondrial oxidative phosphorylation is regulated by fructose 1,6-bisphosphate. A possible role in Crabtree effect induction? J Biol Chem, 2008. 283(40): p. 26948–55.

18. Holden, H.M., I. Rayment, and J.B. Thoden, Structure and function of enzymes of the Leloir pathway for galactose metabolism. J Biol Chem, 2003. 278(45): p. 43885–8.

19. Rempel, A., et al., Glucose catabolism in cancer cells: amplification of the gene encoding type II hexokinase. Cancer Res, 1996. 56(11): p. 2468–71.

20. Wilson, J.E., Isozymes of mammalian hexokinase: structure, subcellular localization and metabolic function. Journal of Experimental Biology, 2003. 206(12): p. 2049–2057.

21. Timson, D.J. and R.J. Reece, Functional analysis of disease-causing mutations in human galactokinase. European Journal of Biochemistry, 2003. 270(8): p. 1767–1774.

22. Sullivan, M.R., et al., Quantification of microenvironmental metabolites in murine cancers reveals determinants of tumor nutrient availability. eLife, 2019. 8: p. e44235.

23. Reinfeld, B.I., et al., Cell-programmed nutrient partitioning in the tumour microenvironment. Nature, 2021. 593(7858): p. 282–288.

24. Meiser, J., et al., Serine one-carbon catabolism with formate overflow. Sci Adv, 2016. 2(10): p. e1601273.

25. Benzarti, M., et al., Metabolic Potential of Cancer Cells in Context of the Metastatic Cascade. Cells, 2020. 9(9).

26. Hyroššová, P., et al., PEPCK-M recoups tumor cell anabolic potential in a PKC-ζ-dependent manner. Cancer Metab, 2021. 9(1): p. 1.

27. Keshet, R., et al., Targeting purine synthesis in ASS1-expressing tumors enhances the response to immune checkpoint inhibitors. Nature Cancer, 2020. 1(9): p. 894–908.

28. Leithner, K., et al., PCK2 activation mediates an adaptive response to glucose depletion in lung cancer. Oncogene, 2015. 34(8): p. 1044–50.

29. Vincent, E.E., et al., Mitochondrial Phosphoenolpyruvate Carboxykinase Regulates Metabolic Adaptation and Enables Glucose-Independent Tumor Growth. Mol Cell, 2015. 60(2): p. 195–207.

30. van der Reest, J., et al., Proteome-wide analysis of cysteine oxidation reveals metabolic sensitivity to redox stress. Nature Communications, 2018. 9(1): p. 1581.

31. Yang, C., et al., Glutamine oxidation maintains the TCA cycle and cell survival during impaired mitochondrial pyruvate transport. Mol Cell, 2014. 56(3): p. 414–424.

32. Lyssiotis, C.A. and A.C. Kimmelman, Metabolic Interactions in the Tumor Microenvironment. Trends in Cell Biology, 2017. 27(11): p. 863–875.

33. Hui, S., et al., Glucose feeds the TCA cycle via circulating lactate. Nature, 2017. 551(7678): p. 115–118.

34. Faubert, B., et al., Lactate Metabolism in Human Lung Tumors. Cell, 2017. 171(2): p. 358–371.e9.

35. Cheung, S.M., et al., Lactate concentration in breast cancer using advanced magnetic resonance spectroscopy. British Journal of Cancer, 2020. 123(2): p. 261–267.

36. Leithner, K., et al., The glycerol backbone of phospholipids derives from noncarbohydrate precursors in starved lung cancer cells. Proceedings of the National Academy of Sciences, 2018. 115(24): p. 6225–6230.

37. Buescher, J.M., et al., A roadmap for interpreting (13)C metabolite labeling patterns from cells. Curr Opin Biotechnol, 2015. 34: p. 189–201.

38. Vande Voorde, J., et al., Improving the metabolic fidelity of cancer models with a physiological cell culture medium. Sci Adv, 2019. 5(1): p. eaau7314.

39. Lemaigre, F.P. and G.G. Rousseau, Transcriptional control of genes that regulate glycolysis and gluconeogenesis in adult liver. Biochem J, 1994. 303 ( Pt 1)(Pt 1): p. 1–14.

40. Kano, A., T. Iwasaki, and M. Shindo, Bongkrekic acid facilitates glycolysis in cultured cells and induces cell death under low glucose conditions. Biochemistry and Biophysics Reports, 2019. 20: p. 100683.

41. Anastasiou, D., et al., Pyruvate kinase M2 activators promote tetramer formation and suppress tumorigenesis. Nat Chem Biol, 2012. 8(10): p. 839–47.

42. Dayton, T.L., T. Jacks, and M.G. Vander Heiden, PKM2, cancer metabolism, and the road ahead. EMBO Rep, 2016. 17(12): p. 1721–1730.

43. Dombrauckas, J.D., B.D. Santarsiero, and A.D. Mesecar, Structural basis for tumor pyruvate kinase M2 allosteric regulation and catalysis. Biochemistry, 2005. 44(27): p. 9417–29.

44. Weinstabl, H., et al., Intracellular Trapping of the Selective Phosphoglycerate Dehydrogenase (PHGDH) Inhibitor BI-4924 Disrupts Serine Biosynthesis. J Med Chem, 2019. 62(17): p. 7976–7997.

45. Smith, G.K., et al., Activity of an NAD-dependent 5,10-methylenetetrahydrofolate dehydrogenase in normal tissue, neoplastic cells, and oncogene-transformed cells. Arch Biochem Biophys, 1990. 283(2): p. 367–71.

46. Lorenz, N.I., et al., AMP-kinase mediates adaptation of glioblastoma cells to conditions of the tumour microenvironment. bioRxiv, 2022: p. 2022.03.25.485624.

47. Fan, J., et al., Quantitative flux analysis reveals folate-dependent NADPH production. Nature, 2014. 510(7504): p. 298–302.

48. Bichi, R., et al., Human chronic lymphocytic leukemia modeled in mouse by targeted TCL1 expression. Proc Natl Acad Sci U S A, 2002. 99(10): p. 6955–60.

49. Chakraborty, S., et al., B-cell receptor signaling and genetic lesions in TP53 and CDKN2A/CDKN2B cooperate in Richter transformation. Blood, 2021. 138(12): p. 1053–1066.

50. Nelson, D.L.a., Lehninger principles of biochemistry. 2017, New York, NY, USA: W.H. Freeman and Company; Houndmills, Basingstoke: Macmillan Higher Education.

51. Chaneton, B., et al., Serine is a natural ligand and allosteric activator of pyruvate kinase M2. Nature, 2012. 491(7424): p. 458–462.

52. Ye, J., et al., Pyruvate kinase M2 promotes de novo serine synthesis to sustain mTORC1 activity and cell proliferation. Proceedings of the National Academy of Sciences, 2012. 109(18): p. 6904–6909.

53. Rossi, M., et al., PHGDH heterogeneity potentiates cancer cell dissemination and metastasis. Nature, 2022. 605(7911): p. 747–753.

54. Muir, A. and M.G. Vander Heiden, The nutrient environment affects therapy. Science, 2018. 360(6392): p. 962–963.

55. Zheng, Y., et al., Mitochondrial One-Carbon Pathway Supports Cytosolic Folate Integrity in Cancer Cells. Cell, 2018. 175(6): p. 1546–1560.e17.

56. Kiweler, N., et al., Mitochondria preserve an autarkic one-carbon cycle to confer growth-independent cancer cell migration and metastasis. Nat Commun, 2022. 13(1): p. 2699.

57. Hiller, K., et al., MetaboliteDetector: comprehensive analysis tool for targeted and nontargeted GC/MS based metabolome analysis. Anal Chem, 2009. 81(9): p. 3429–39.

58. Wang, T., et al., Inosine is an alternative carbon source for CD8(+)-T-cell function under glucose restriction. Nat Metab, 2020. 2(7): p. 635–647.

59. Dranka, B.P., et al., Assessing bioenergetic function in response to oxidative stress by metabolic profiling. Free Radic Biol Med, 2011. 51(9): p. 1621–35.

